# A meta-analysis of environmental sequencing data reveals the global distribution and hidden diversity of marine anaerobic ciliates

**DOI:** 10.64898/2025.12.15.694440

**Authors:** Anna Schrecengost, Erin Frates, Alia Al-Haj, Robinson W. Fulweiler, Roxanne Beinart

## Abstract

Anaerobic protists are diverse, ecologically important members of anoxic microbial communities, acting as grazers, nutrient cyclers, and partners in multi-domain associations, yet remain understudied relative to anaerobic prokaryotes. Ciliates are particularly abundant and diverse in anoxia, but their global diversity and distribution are largely unknown. Here, we conducted a meta-analysis of public 18S rDNA datasets, along with one dataset generated here, to assess the global diversity and ecology of marine anaerobic ciliates. Using a novel pipeline, we processed 2854 samples from 42 studies spanning 19 habitat types. We recovered 3196 anaerobic ciliate amplicon sequence variants (ASVs) across all described lineages. Based on clade-specific divergence thresholds derived from phylogenetic distances, 28.4-46.3% of ASVs qualified as novel. Most sequences belonged to the poorly described plagiopylean family Epalxellidae, suggesting a large reservoir of undescribed diversity in this clade. Community comparisons revealed close phylogenetic similarities between some shallow-water and deep-sea assemblages, suggesting that shared redox conditions may shape communities more than water depth. Our results demonstrate that marine anaerobic ciliates are globally distributed, taxonomically diverse, and rich in novel lineages. This study provides a framework for leveraging environmental sequencing data to better understand the diversity and ecology of neglected protist lineages and under-sampled habitats.

## INTRODUCTION

Protists are among the most common and abundant organisms on the planet – some estimate that they may constitute over 85% of the overall eukaryotic diversity in the sunlit ocean, with similarly high proportions likely across other environments (1). These single-celled eukaryotes play essential ecological roles, particularly in aquatic environments: they constitute a major proportion of microbial activity and biomass, cycle essential nutrients, and are major contributors to the biological carbon pump which sequesters approximately 2.5 Pg carbon per year (2–6). Phototrophic protists form the foundation of aquatic food webs, while heterotrophic protists are essential members of microbial loops and major contributors to mortality. Most, if not all, of these processes are involved in the delicate biogeochemical feedback loops which regulate our rapidly changing climate, making it urgent to characterize this “unseen majority”. Specifically, by understanding the diversity of anaerobic protists and the factors that influence their community composition, we can begin to understand how they will affect and be affected by climate change (7). In the past several decades, advancements in DNA sequencing and metabarcoding have greatly expanded our ability to study previously elusive protistan species and allowed us to explore protistan communities from previously inaccessible or largely unexplored environments, including oxygen-depleted marine environments (8,9). Most metabarcoding studies of marine protists are small-scale and limited in geographic scope, capturing a diversity of habitats but often employing different methods to do so, which complicates our ability to coherently synthesize this information. Extensive global sampling efforts focused on planktonic species (1,10,11), with relatively few targeting benthic or anaerobic taxa (5).

While most eukaryotic life relies on oxygen for respiration, a huge diversity of protists live in anoxia, and ciliates are particularly active, large, and diverse members of these understudied communities. They are a diverse and versatile group of protists known for their ability to establish various symbiotic partnerships (12–14) and inhabit a wide array of environments, including oxygen-depleted habitats such as aquatic sediments, the hypoxic water column, and the digestive tracts of animals (15–19). Most ciliate lineages include representatives adapted to some degree of anoxia, ranging from microaerophilic to obligately anaerobic (20,21). Indeed, analyses of phylogenetic and genomic data have made it clear that ciliates have transitioned from an aerobic to anaerobic lifestyle multiple times throughout their evolution (15,22,23). Whereas anaerobic prokaryotes are relatively well-studied and appreciated for their large impact on biogeochemical cycles and greenhouse gas production (24,25), microbial eukaryotes have been largely neglected in our accounting of anaerobic microbial communities. Anaerobic eukaryotes originating from marine environments have been especially overlooked, as most of the described species originate from freshwater habitats (20). As hypoxic zones of the ocean have been expanding and the concentration of dissolved oxygen in aquatic habitats has been continuously decreasing over the past several decades (26,27) a full accounting of and appreciation for the various members of marine microbial communities especially grows in importance.

Given that much of our current understanding of anaerobic ciliate diversity stems from morphological surveys without any known DNA sequences (28) or from a few cultured representatives, metabarcoding is a promising, though underused, approach to uncover novel lineages and assess the full diversity and distribution of protist groups. Molecular studies of oxygen-depleted marine environments have been crucial in our understanding of the genetic diversity of anaerobic ciliates, generating the majority of the sequences which our reference phylogenies and taxonomic databases are based on (17,29–34) and uncovering novel lineages, some of which are thus far understood to be endemic to specific locations (35). In terms of metabarcoding studies, there have been various geographically limited studies targeting protists from anoxic environments (36–38) which used older pyrosequencing technologies, as well as newer, Illumina sequencing studies of benthic protists in intertidal (39–41), hydrothermal vent (42,43), methane seep (44), and deep sea (45) environments, and of protists in bottom waters adjacent to hydrothermal vents (46,47). Many of these studies reported anaerobic ciliate taxa. However, thus far studies targeting microbial eukaryotes living in oxygen-depleted environments have been localized and no efforts have been made to synthesize this wealth of sequencing data in a meta-analysis.

A comprehensive examination of existing marine anaerobic ciliate sequencing data would allow us to gain a better understanding of their global distribution and overall diversity (our current knowledge of which is primarily limited to local, culture-based studies) as well as assess the potential for novel lineages. Previously, we used culture–based approaches to survey marine anaerobic ciliates from intertidal sediments in Southern New England, and recovered a diversity of *Plagiopyla* and *Metopus* species, some of which were novel (48–50). In this study, we generated 18S rDNA amplicon sequencing data isolated from subtidal sediments in Narragansett Bay, Rhode Island. Additionally, we mined 41 publicly available, environmental 18S rDNA datasets originating from marine, oxygen-depleted environments for the presence of anaerobic ciliate sequences. We did this in a systematic manner via the creation of a Snakemake pipeline which takes raw reads from high-throughput sequencing (HTS) datasets as input and outputs per-sample amplicon sequence variants (ASVs) and per-study metadata. We then used multilevel phylogenetic placement (51,52) to identify and taxonomically assign ciliates from three major anaerobic ciliate clades: the APM clade (15), Plagiopylea (53), and Anaerocyclidiidae (54). These groups are the most commonly recovered in both culture-based and independent studies, and only a handful of sequences outside of these groups are known (Table S3). Phylogenetic placement has been shown to outperform traditional pairwise alignment-based taxonomic assignments in other ciliate groups (e.g. colpodean ciliates) (55). It is a promising method particularly for groups which are underrepresented in reference databases, e.g., anaerobic ciliates, because it incorporates phylogenetic information and can retain deeply divergent sequences which would not be assigned with traditional methods. Given the frequency of discovery of novel lineages of anaerobic ciliates with low-throughput methods (15,35,54), divergent sequences are likely abundant in these underexplored environments. Additionally, it enables us to analyze studies which used different primers or even sequenced different regions of the 18S rRNA gene together, is more robust to very short reads, and can recover novel taxa (56).

The goal of this study was to use publicly available data to explore anaerobic ciliate diversity across as many marine oxygen-depleted habitats as possible, including those in which they are not well-characterized, in order to explore their total global diversity, understand more about their distribution across depths and among habitat types, and identify potential novel lineages. By synthesizing this global dataset of phylogenetically-placed anaerobic ciliate sequences, we recovered a diversity of sequences across all free-living anaerobic ciliate families from habitats as disparate as hydrothermal vents to intertidal sediment. The vast majority of the sequences that we recovered across all habitats belonged to the little-known and poorly-described plagiopylean family Epalxellidae, which thus far only includes uncultured representatives from a diversity of habitats (57,58), suggesting a substantial reservoir of uncharacterized diversity and the potential for future exploration of this group. We also found that, while anaerobic ciliate communities were largely distinct across water and sediment, they did not appear to be structured by water depth, with some deep-sea communities resembling those in shallow coastal settings. These results have implications not only for our understanding of anaerobic ciliate diversity but also provide us with more information about which habitats and groups should be prioritized for more robust sampling and illustrate the power of a meta-analysis approach for understudied and – sampled protist groups. The pipeline presented here provides a framework that other researchers can apply to explore the global diversity of their protist groups or habitats of interest. This approach holds promise for advancing our understanding of neglected microbial eukaryotes, particularly those from extreme, inaccessible habitats, by leveraging the wealth of existing sequencing data deposited in public repositories.

## METHODS

### Study selection

We performed a literature review to identify 18S rRNA gene high-throughput sequencing datasets which might possibly contain marine anaerobic species and which were sequenced with Illumina technology. We searched Google Scholar with the following search terms: “marine sediment”, “intertidal”, “18S”, “Illumina” “microeukaryote”, “ciliate”, “protist”, “anoxic”, “oxygen-depleted”, “hydrothermal vent”, “oxygen-minimum zone”, “cold seep”, “methane seep”. We also searched for 18S amplicon studies which reported the presence of anaerobic ciliate taxa. We only included peer-reviewed studies which were written in English. Additional studies were sometimes obtained through the papers which were referenced in the studies from our search results. This literature review was completed in January of 2024, and no studies published after that date were included in the analysis. Additionally, we included one new study of sediment 18S rRNA gene sequences that we generated from Narragansett Bay, Rhode Island (USA). All studies included in our analysis are listed and described in Table S1 and more information about data generation, study selection, and metadata is available in the Supplemental Methods.

### SRA to ASV Snakemake pipeline: Raw data retrieval, processing into per-sample ASVs, and taxonomic assignment

Raw data retrieval and processing into ASVs was carried out via QIIME2 and Snakemake (Figure S1). All steps were implemented as modular Snakemake rules and executed in a reproducible environment managed by Conda. Primer removal, merging of paired end reads, and denoising were performed in QIIME2 using cutadapt, vsearch, and deblur, respectively (59–62). Taxonomic assignment of the resulting representative sequences was performed using a Naive Bayes classifier trained on the PR^2^ reference database (63). Classification output was used for downstream filtering, resulting in per-study count tables for sequences assigned to Cilophora as well as Unassigned sequences. After all studies were run through the Snakemake pipeline, Ciliophora-assigned and Unassigned ASVs and ASV count tables, respectively, were merged for downstream processing.

### Multi-level phylogenetic placement of ASVs onto reference eukaryotic, ciliate, and anaerobic ciliate clade phylogenetic trees

The Ciliophora-assigned and Unassigned ASVs were phylogenetically placed onto a series of reference trees in order to identify and extract anaerobic ciliate sequences. Previously published eukaryotic and ciliate reference trees which were designed for the phylogenetic placement of short reads were used for this purpose (52),(64) (Figure 2). Anaerobic ciliate clade trees were constructed for the placement of per-clade ASVs (details available in the Supplemental Methods). For the phylogenetic placements, we followed a multi-level phylogenetic placement strategy as described elsewhere (51,52). ASVs which were unassigned with QIIME2 taxonomic assignment were phylogenetically placed onto the Eukaryota reference tree. Ciliate sequences were extracted from those placements, combined with the sequences assigned to Ciliophora with QIIME2, and then placed onto the Ciliophora reference tree. Sequences assigned to each anaerobic ciliate clade were extracted from these Ciliophora placements and placed onto their respective reference trees. In the case of the APM clade, sequences which were assigned to Armophorea on the Ciliophora tree included all APM lineages, following the taxonomic scheme described in Ratjer and Dunthorn (2021) (64).

Phylogenetic placement was conducted as follows: ASVs were aligned to the full-length reference multiple sequence alignment using PaPaRa v2.5 (65), which performs phylogeny-aware alignment extension with the reference phylogenetic tree, reference alignment, and query sequences (ASVs) as input. This alignment was split into reference and query subsets with the epa-ng –-split command (66). Maximum likelihood model evaluation for the reference alignment was performed using the raxml-ng –-evaluate command, which identifies the best-fitting model parameters for placement (67). Phylogenetic placement of query sequences onto the reference tree was conducted using EPA-NG (66), with filtering parameters to retain the top 100 placements per query with a minimum likelihood weight ratio (LWR) of 0.99. This produced a .jplace file containing the distribution of placements for each query across the reference tree.

### Analysis of phylogenetic placement results and anaerobic ciliate sequences

Gappa (68) was used to analyze and visualize phylogenetic placement results. High-confidence phylogenetic placements of anaerobic ciliate ASVs were filtered based on placement uncertainty (EDPL ≤ 0.05). Taxonomic assignments were generated from these placements using the command gappa examine assign.

For each reference tree, phylogenetic placements were visualized with the Interactive Tree of Life (iTOL v7) (69)). The global distribution of sampling locations was visualized by plotting the geographic coordinates from sample metadata using the sf and ggplot2 packages in R (70,71). ASV relative abundance as well as agglomerated taxonomic presence/absence were calculated per sample, and fractional taxonomic abundance plots were created with the PHYLOSEQ package in R (72,73). Low abundance reads (<2) for each sample were removed prior to analysis.

The phylogenetic divergence between anaerobic ciliate communities from different habitat types was assessed using both the Kantorovich-Rubinstein Distance (KRD) and squash clustering. Prior to analysis, .jplace files were split per sample and grouped by habitat type. The KRD is a generalization of Unifrac distance which calculates the minimum effort needed to transform one community’s placement distribution into another’s by moving mass along the tree (74). Squash clustering is a hierarchical clustering method designed for phylogenetic placement data (75).

For each anaerobic ciliate clade, per-clade phylogenetic trees were re-inferred by re-aligning clade-assigned ASVs with reference sequences. Following tree inference, we assessed the phylogenetic distances between ASVs and references by computing the minimum patristic distance between each query and the reference sequences, yielding a per-query estimate of evolutionary divergence from the closest known relative. These distances were visualized using ridge plots in R to compare clade-specific patterns of phylogenetic novelty. Percent identity to the closest reference sequence for each query was also determined via BLASTn against the PR^2^ database.

ASVs were categorized as “novel” sequences based on sequence divergence as calculated via patristic distances. For each anaerobic ciliate clade (as shown in Figures 4 and 5), the average and maximum patristic distance between reference sequences was calculated, and these were used as thresholds. For each clade, ASVs which had greater patristic distances to their closest neighbor in the reference tree than these thresholds were classified as novel (Table S4).

## RESULTS AND DISCUSSION

### Recovery of a diversity of free-living anaerobic ciliate sequences from global marine environmental datasets

In total, we processed 2854 samples from 43 studies representing 19 distinct marine oxygen-depleted habitats (Figure 1, Table S2). We recovered 36,358 ciliate-assigned ASVs in 2030 of those samples, and 3196 anaerobic ciliate ASVs across 649 samples. Overall, we were able to recover a diversity of anaerobic ciliates from all habitats surveyed here, except from mangrove sediment and bathypelagic water (Figures 1, 3). We found anaerobic ciliates from every free-living marine genus within the clades included in this study (APM, Plagiopylea, and Anaerocycliididae), as well as many sequences which were unrelated to known genera (discussed below). Additionally, we recovered ciliate sequences from across virtually the entire diversity of Ciliophora (Figure 2). Although these aerobic ciliate sequences are not investigated here, our recovery of them may reflect both the remarkable ecological flexibility of ciliates, as well as the diversity of anoxic marine habitats, which can exist in microniches both spatially and temporally, e.g., in hydrothermal vent fluid, associated with surfaces, and even in sediment (19,76,77). Assuming that these sequences originated from live ciliates, it is possible that the samples included in this analysis are mismatched with the microniches where anoxia generally exists, and that many, if not the majority, of samples included originated from primarily oxic or micro-oxic habitats.

**Figure 1.**
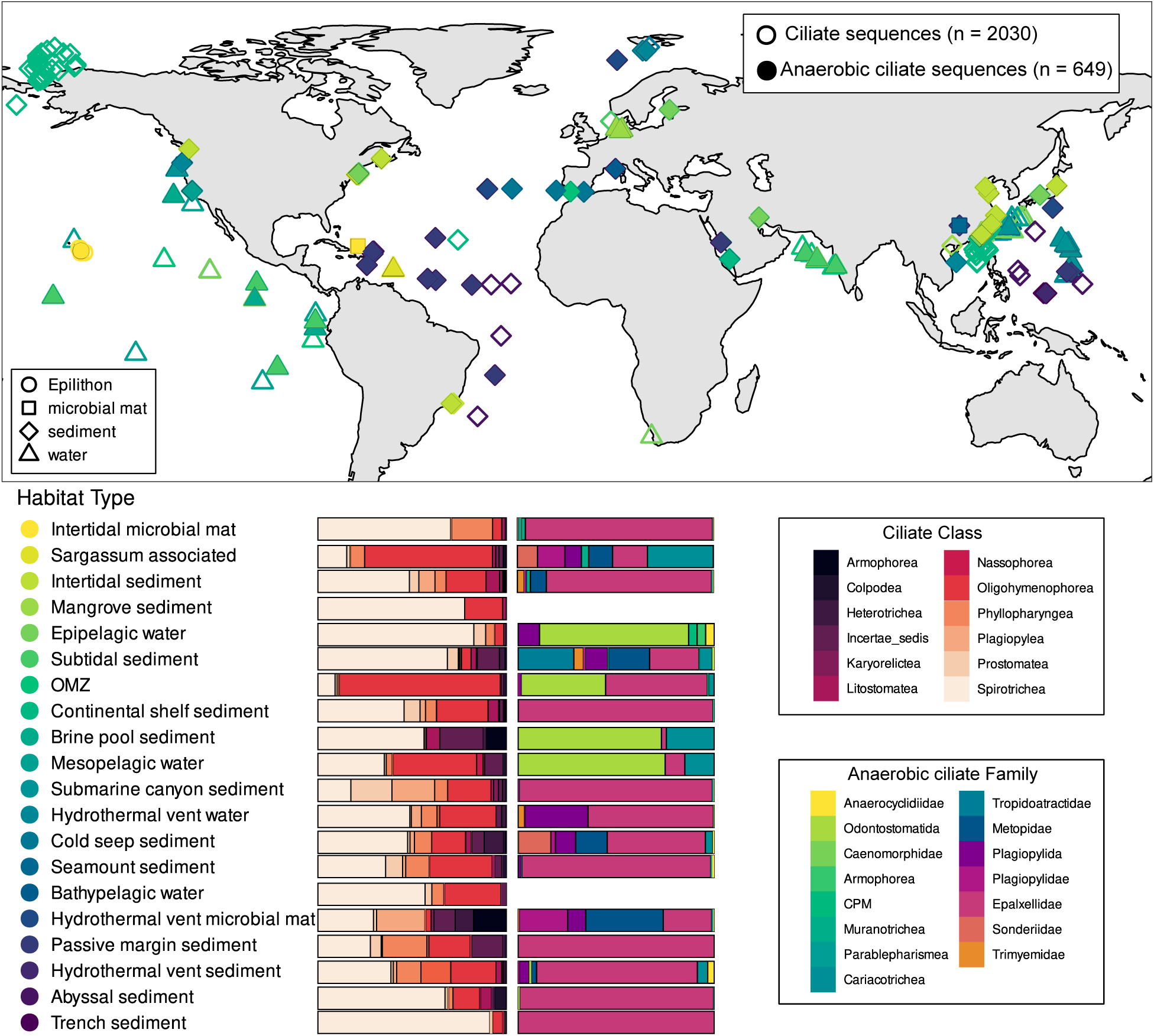
Top: Map showing location of samples included in this study which contained ciliate sequences (empty outlined points) and anaerobic ciliate sequences (filled in points). Points are colored by habitat type, which is described in the legend on the bottom, and substrate type, which is described in the legend within the map. Bottom: compositional bar plots describing the relative abundance of different ciliate classes (left) and a anaerobic ciliate families (right) recovered in the samples, separated by habitat type.

**Figure 2:**
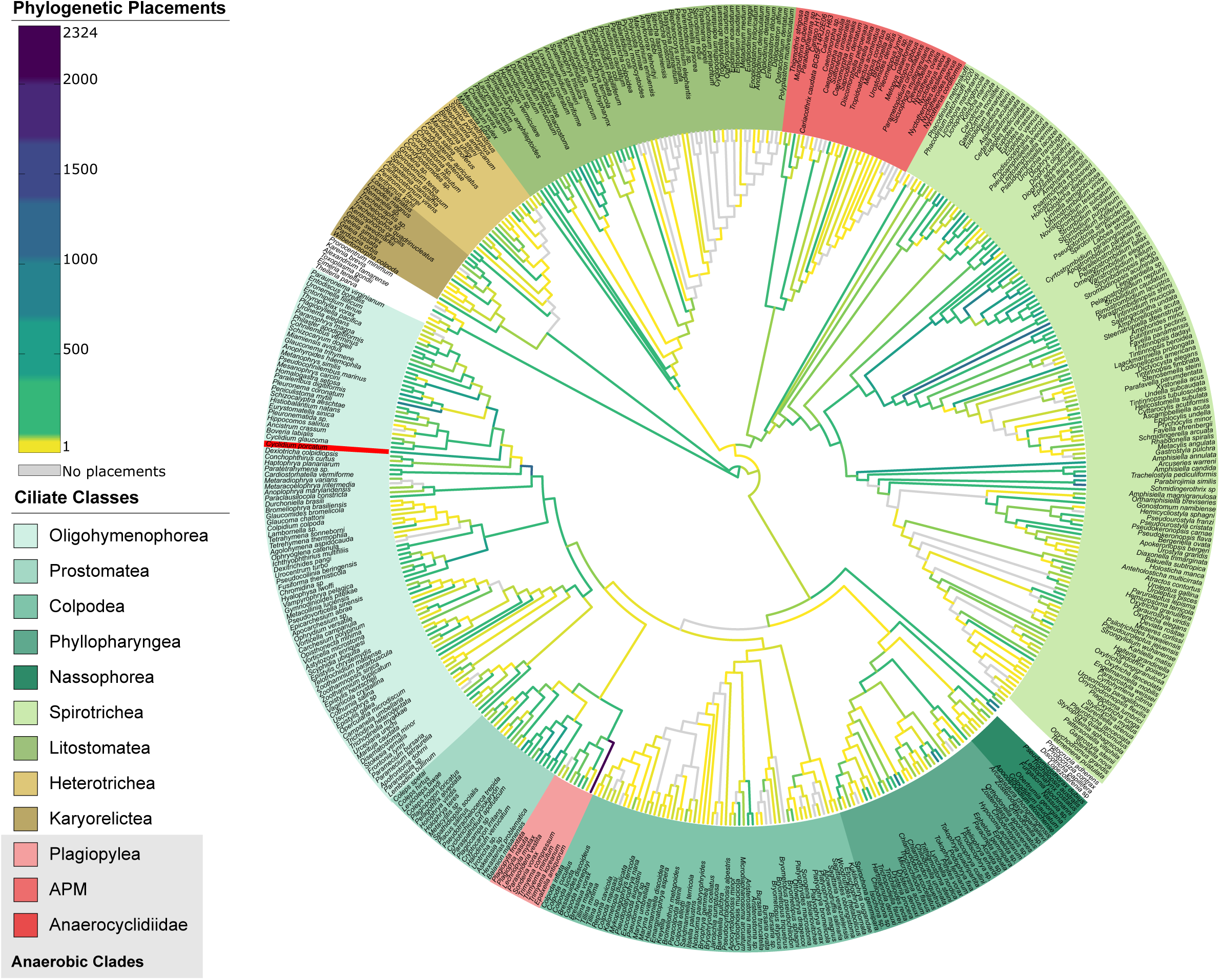
Phylogenetic placement of pre-filtered ASVs onto Ciliophora reference tree. A full length 18S rRNA gene reference phylogenetic tree (69) was used in the phylogenetic placement of pre-filtered ASVs, which were either assigned to Ciliophora in QIIME2 or were unassasigned in QIIME2 but later assigned to Ciliophora via phylogenetic placement onto a reference Eukaryota phylogenetic tree. Ciliate classes are colored at the tips, and anaerobic clades are colored in red. Color on branches indicates the distribution of top phylogenetic placements for each query, which corresponds to their phylogenetic placement-based taxonomic assignment.

The only habitats in which we recovered ciliate reads but no anaerobic ciliate reads were mangrove sediment and bathypelagic water; although, in the case of mangrove habitats, the number samples included (20 samples) and number of total ciliate reads recovered were very low (104 ASVs). Indeed, mangrove sediments do harbor anaerobic protists (48,78) but global environmental sequencing data from mangroves and tropical estuaries is lacking (79). Therefore, it is likely an issue of undersampling rather than true absence. Indeed, the compositional nature inherent to HTS data means that we can never truly know what zeros represent in this data. The issues inherent in microbiome data analysis, including potential biases at each step such as sample acquisition and processing (80), are multiplied in the case of a meta-analysis such as this one, where each study carried out different sampling strategies, used different primer sets, and underwent different sequencing efforts. Additionally, protist spatial and temporal distribution is heterogeneous, and anaerobic protists are traditionally thought to be rare, although it is likely they are not as rare as once thought (9,81). The bioinformatic methodology in this study was designed to minimize differences between datasets – when possible, we used the same parameters and trimmed reads to the exact same regions to minimize biases. Additionally, we used the deblur algorithm for denoising, which is the most conservative of the denoising algorithms and was designed for use in meta-analyses (60). We also used a phylogenetic placement method, which allowed us to analyze all ASVs together regardless of primer set and place the sequences into a phylogenetic context (56).

### Anaerobic ciliate communities appear similar across some deep sea and shallow water habitats

To compare anaerobic ciliate communities across different global marine habitat types, we performed phylogenetic placement-based beta diversity analyses (Figures 3 and S5). From these results, we observed that water column samples from oxygen-minimum zones, anoxic basins, estuarine water, etc. are more similar to each other than to other samples. We also observed close similarity between anaerobic ciliate communities from hydrothermal vents, intertidal zones, and cold seep sediments. Microbial mat and sediment samples from hydrothermal and intertidal/subtidal environments consistently grouped together. Specifically, hydrothermal vent microbial mat and subtidal sediment were most similar, as were hydrothermal vent sediment and intertidal sediment and deep cold seep sediment and intertidal microbial mat. Hydrothermal vent fluid and hydrothermal vent fluid influenced seawater community compositions were most similar as well.

**Figure 3.**
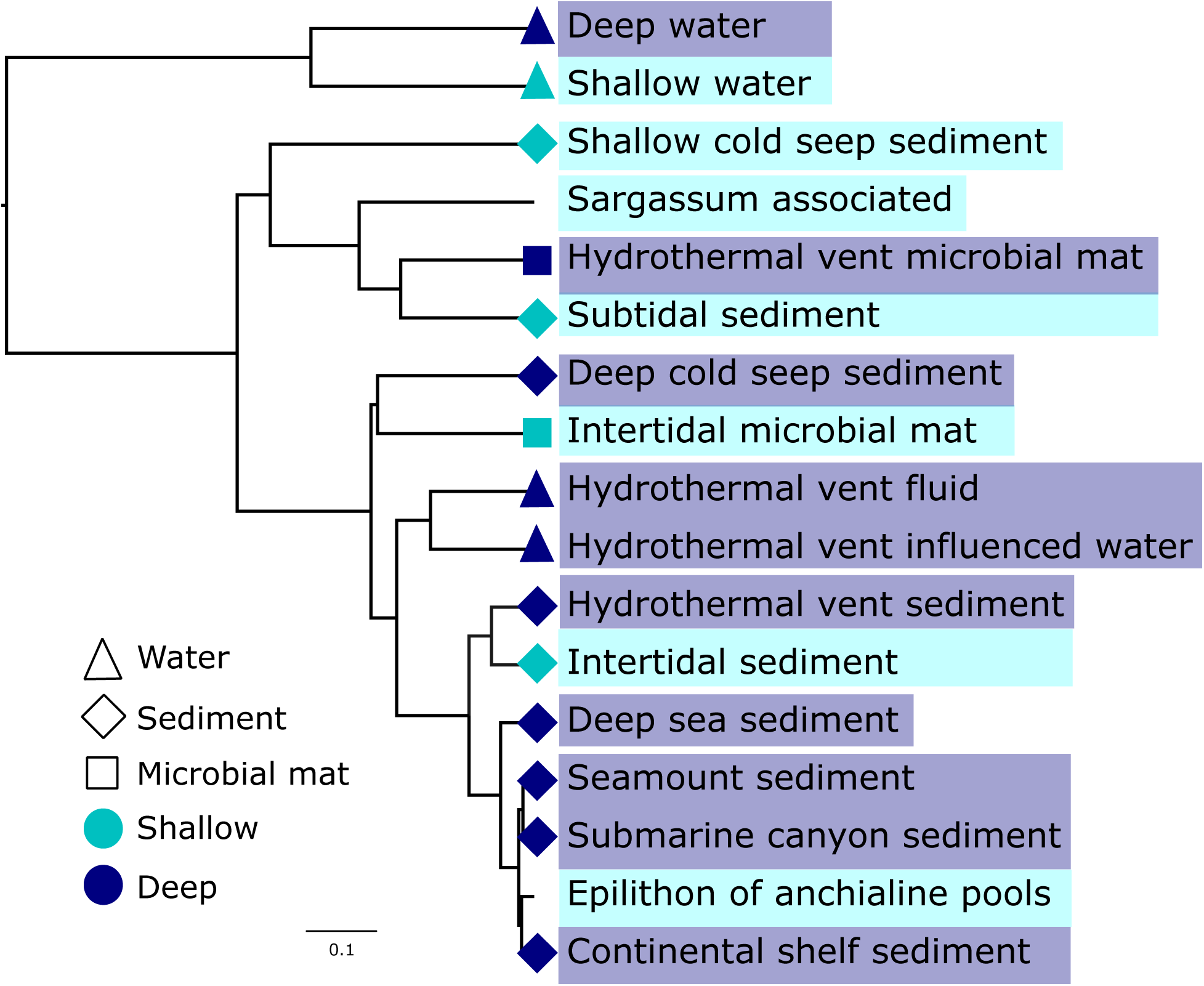
Hierarchical clustering of Kantorovich-Rubinstein Distance (KRD) between samples grouped by habitat type. Branch tips are colored by water depth, with deep defined as greater than 300 m, and branch node shapes indicate sample substrate.

Presence/absence analysis of anaerobic ciliate taxa across these same habitat types also suggests that deep-sea cold seep and hydrothermal vent and shallow water intertidal and subtidal habitats share similar community compositions (Figure 4). These habitats were comprised of the most samples which contain these taxa. Most habitats were dominated by the presence of Epalxellidae, with Plagiopylida, Cariacotrichea, and Anaerocyclidiidae also very prevalent across many habitats. Interestingly, Sonderiidae ciliates were only recovered from intertidal sediment, subtidal sediment, cold seep sediment, and hydrothermal vent microbial mat, as well as Sargassum-associated samples. On the other hand, water column habitats lacked as many detectable anaerobic ciliate taxa as sediment and microbial mat habitats.

**Figure 4.**
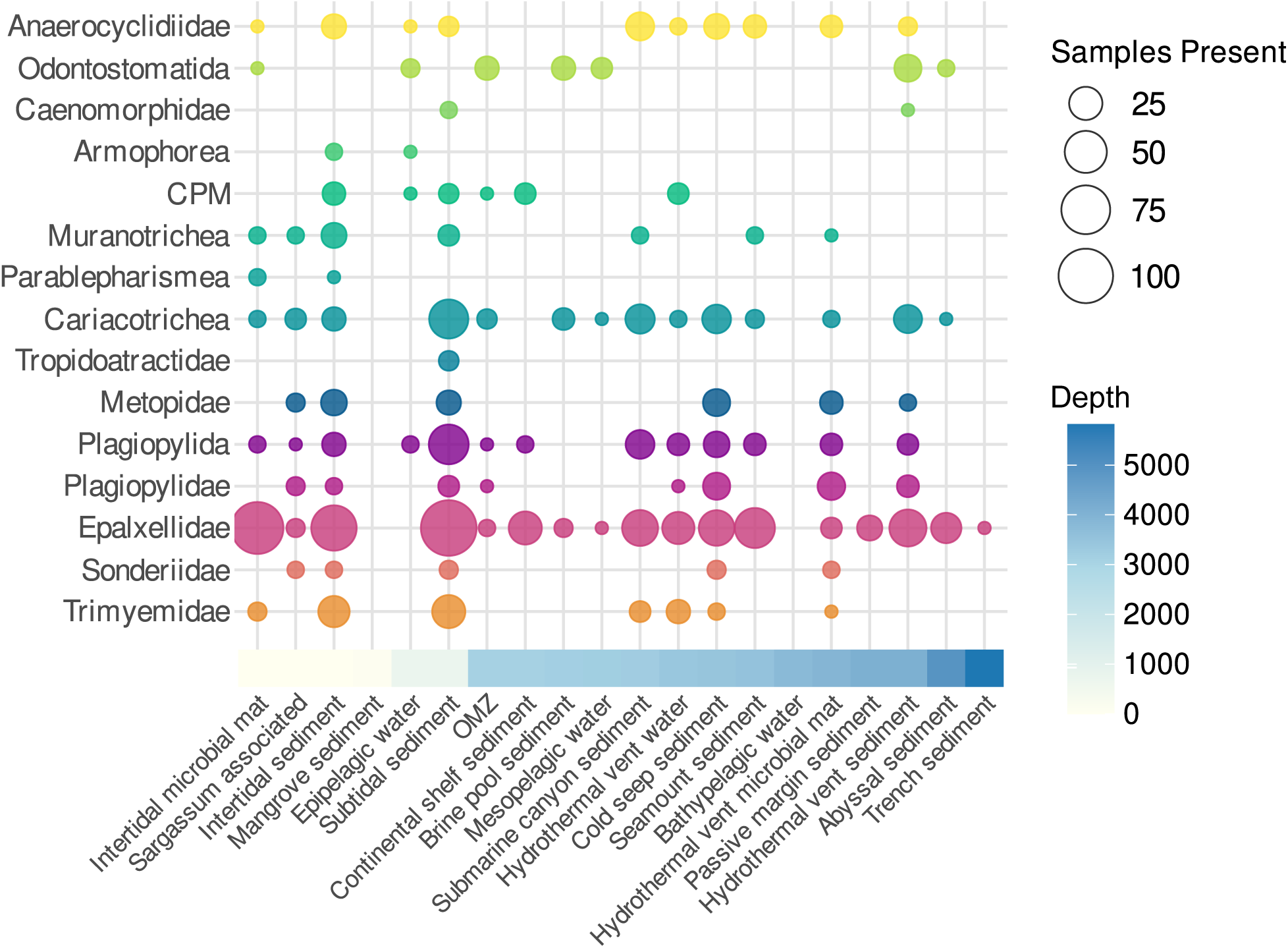
Presence-absence bubble plot illustrating prevalence of major anaerobic ciliate taxa across marine oxygen-depleted habitats. Habitat types are organized and colored by average depth. Bubble sizes correspond to the number of samples that the taxa was present in for each habitat.

These clustering patterns may suggest similarities in anaerobic ciliate community composition across deep-sea and shallow-water habitats that may be due to biogeochemical similarities in these habitats, despite dramatic differences in depth/pressure. Hydrothermal vent fluid samples clustered with sediment and microbial mat samples, while other water samples (e.g., OMZ) did not. This may be due to the chemical similarity of vent fluid and sediment – vent fluids contain high concentrations of reduced chemicals, such as hydrogen sulfide, methane, and hydrogen, etc. (82); similarly, sediments contain redox gradients where these same chemicals dominate as electron acceptors at various depth horizons, and these chemicals fuel anaerobic metabolisms (83). Oxygen minimum zones and oxygen deficient water columns more generally have various degrees of oxygen depletion. Most are not fully anoxic and do not have appreciable concentrations of sulfides or other chemical reductants (84). Additionally, the clustering observed between shallow-water microbial mats and deep-sea sediments (e.g., at cold seeps and hydrothermal vents, Figures 3 and S5) is possibly influenced by their chemical and structural similarity. All of these habitats are rich in organic matter and are similarly physically structured by vertical redox gradients (85). It is also possible that these patterns in anaerobic ciliate community composition are influenced by similarities in the bacterial and archaeal communities across these habitats and driven by interdomain biological interactions between these groups. For example, Beggiatoa microbial mats are found globally across shallow to deep water in coastal, mangrove, hydrothermal vent, and cold seep sediments, among other habitats that interface the underlying sulfidic and overlying oxic zones (86). Anaerobic ciliates interact with bacteria and archaea via grazing as well as symbioses that they have established with methanogenic archaea, sulfate reducing bacteria, and other bacteria (20,87), and therefore similarities in bacterial and archaeal communities across these deep-sea and shallow water habitats could also influence similarities in anaerobic ciliate community composition.

Interestingly, multiple previous studies have demonstrated that protist communities are significantly different across depth in the marine water column (88,89) and across overlying water column depth in the sediment (90,91). Depth has been shown to greatly influence the composition of benthic ciliate communities (45) but not always pelagic communities (92). In both deep-sea and shallow habitats, benthic protists have been found to be shaped more by depth than geography, where even geographically disparate communities are highly similar at the same depth profile (91,93). And yet, our results indicate that anaerobic ciliate communities in some shallow habitats (e.g. intertidal sediments) are very similar to those in some deep-sea habitats (e.g., hydrothermal vent sediment). In the aforementioned cases, it is possible that depth is a proxy for many other environmental factors, such as light availability, pressure, salinity, and temperature. [AA1] In anoxic sediment and microbial mat habitats, many of these environmental factors are similar: they have similar redox conditions, no light availability, high microbial biomass and organic matter loading, and are rich in reduced compounds such as hydrogen sulfide. Our results suggest that these factors are what influence anaerobic ciliate community composition rather than depth or geographic distance. Indeed, identical anaerobic ciliate species have been isolated from similar habitats across the globe, and, in some cases, they even host identical archaeal symbionts (48,49). While the Baas Becking hypothesis, i.e. “everything is everywhere, but the environment selects”, has been largely disproven for protist communities which have been shown to be biogeographically structured (9), it is possible that this is truer for extremophile protist communities such as anaerobic protists which have exploited niches largely inaccessible to most other microbial eukaryotes (23). Large-scale meta-analyses such as this one help us better understand the global diversity and distribution of different protist groups and how they are influenced by various environmental conditions. However, it is also important to consider that marker genes such as the 18S rRNA gene do not have enough phylogenetic resolution to reliably distinguish species, and so sequences which seem identical may actually be phylogenetically, pheno– or genotypically distinct.

### Recovery of a high prevalence of anaerobic ciliate sequences from the little-known plagiopylean family Epalxellidae and other divergent sequences

One of the most striking results from our analysis is the prevalence of Epalxellidae ciliates across virtually all marine oxygen-depleted habitats surveyed (Figures 2 and 4). Not only was this the most common anaerobic ciliate taxa observed across all the samples (2206 out of 3196 (69%) of all anaerobic ciliate placements were on this clade), but even among all ciliate phylogenetic placements, by far the greatest density of placements was observed on this little-known clade (Figure 2). Additionally, ciliates from this family were found in habitats where no other anaerobes were recovered, such as some deep-sea sediments (Figure 4). Taking a closer look at the Plagiopylea placement results (Figure S2), most of these sequences were placed on the basal branch of Epalxellidae or on/near the *Epalxella antiquorum* branch.

Epalxellidae was originally described as an odontostomatid based on morphology, but subsequent genotyping grouped it with the plagiopyleans (57). It represents a basal lineage within the Plagiopylea and only contains one described, although uncultivated, representative, *Epalxella antiquorum*. The remaining representatives originate from a range of habitats: cold seep sediment, the Cariaco basin, the Black Sea, a benthic diatom film, sewage treatment sludge, an anoxic fjord, as well as a freshwater lake (Figure S2). The freshwater lake representative harbors unique bacterial symbionts which generate ATP for the host via denitrification (94). Combined with the described methanogenic archaeal symbionts of other plagiopylean ciliates and the huge diversity of habitats where it is found globally, this suggests a remarkable ecological diversity within this little-known anaerobic ciliate family. The abundance of ASVs assigned to this ill-defined group as well as its prevalence across the global habitats surveyed here suggests that there is large pool of uncharacterized diversity of ciliates from this family, and future studies should prioritize understanding anaerobic protists and underexplored protist groups in general.

Approximately half of the ASVs assigned to the APM clade (203 out of 291) belonged to a poorly described subgroup within the APM, which we refer to here as the CPM clade (Cariacotrichea, Parablepharismea, and Muranotrichea; Figure S3). All of these ciliate classes are little known, deep-branching, and have only been described a few times in the literature. There is currently only one known species from the class Cariachotrichea, *Cariacothrix caudata*, which has been isolated from the water column of the deep anoxic Cariaco basin water column (95,96). Other sequences which are included in this clade in this and other analyses (97) were isolated from a deep-sea methane seep (44) and in an oxygen minimum zone (98). After Epalxellidae and Plagiopylida ciliates, Cariachotrichea was the most prevalent group in our analysis, recovered from both shallow water and deep sea habitats (Figure 4). Parablepharismea contains two recently sequenced species isolated from brackish lagoon sediment (99), as well as species from salt pond sediment, although those are not included here because they are too similar to the other species (15). Muranotrichea was only recently established as a novel ciliate class and contains two described genera which were isolated from marine and brackish beach, mangrove, and salt pond sediment (15).

Not only did we recover many sequences from these poorly described groups, some of which were assigned to basal branches, but many of these sequences, along with sequences from other anaerobic ciliate groups, were quite unrelated to reference sequences, both based on pairwise nucleotide identity and phylogenetic distance (Figure 5, Table S4). Sequences assigned to Trimyemidae, Sonderiidae, Epalxellidae, Plagipylida, Tropidoatractidae, Cariacotrichea, the CPM clade, Armophorea, Odontostomatida, and Anaerocyclidiidae (or, the majority of groups surveyed here) showed broad and diffuse distributions of pairwise identity with reference sequences, with many closer to 90% identity, which has been used as approximate threshold for sequences within the same family (52). Sequences assigned to Odontostomatida were the most divergent, particularly in regard to phylogenetic distance, although the phylogenetic position of this group is poorly resolved, there are only three known sequences, and it is usually the longest branching group in Ciliophora phylogenies (58); therefore, larger phylogenetic distances are likely unreliable.

**Figure 5.**
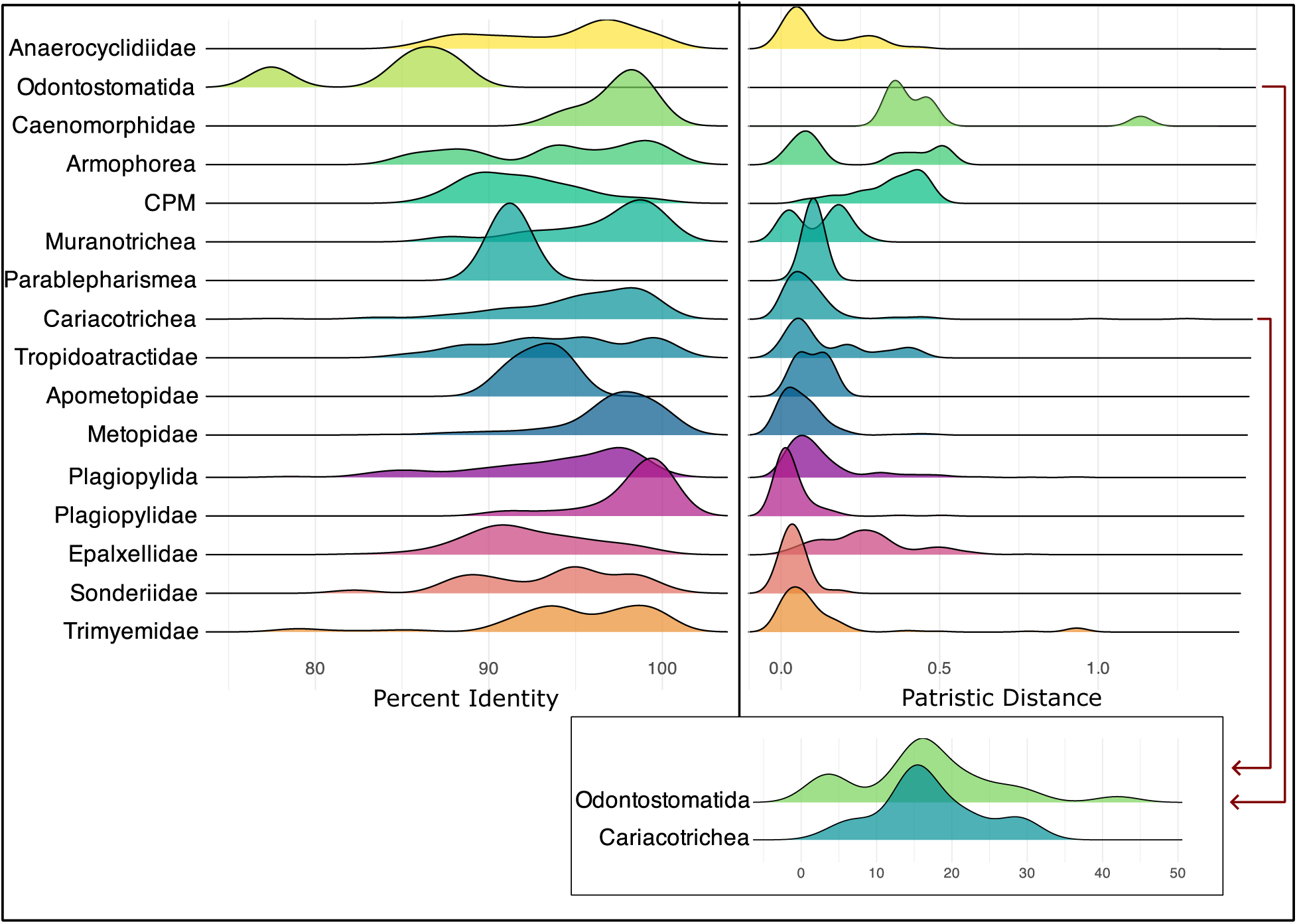
Left: Pairwise percent identity of anaerobic ciliate sequences to their closest reference hit in the PR^2^ database, as determined by BLASTn. Sequences are grouped by taxa and results are shown in a ridge plot. Right: Patristic distance of anaerobic ciliate sequences to the sequence which they are most closely related to phylogenetically, based on a re-inferred phylogenetic tree which includes both reference and ASV sequences. Sequences are grouped by taxa and results are shown in a ridge plot. The patristic distance refers to the sum of the branch lengths between the ASV and the closest reference sequence. Distances larger than 1.5 are excluded from the main plot and instead shown in the inset at the bottom.

We classified ASVs as novel if the phylogenetic distance to their closest neighbor on the reference tree (which was calculated from a phylogenetic tree containing both ASVs and reference sequences) was greater than the average phylogenetic distance among sequences within their assigned clades (Table S4). We also classified ASVs much more conservatively using the maximum per-clade phylogenetic distances as the threshold, and here we report the estimated proportion of novel ASVs per clade as a range. Using this criterion, 31.8-49.6% of the ASVs in Plagiopylea were novel, as were 8.9-32.3% of the ASVs within the APM clade and 0-14.78% of the ASVs in Anaerocyclidiidae. Smaller clades within these broader lineages contained various proportions of novel sequences. For example, 40-57.6% of ASVs classified within Epalxellidae were novel, whereas only 6.7% of ASVs from *Plagiopyla* were novel. We also recovered a large proportion of novel sequences assigned to the CPM clade (0-61.5%). Overall, most of the anaerobic ciliate clades recovered in this study contained substantial proportions of novel ASVs (at least 20%). Taken together, these results suggest that our meta-analysis uncovered a large amount of undescribed diversity and demonstrates the utility of a phylogenetic placement-based meta-analysis approach such as this for the exploration of understudied protist groups and habitats.

## CONCLUSIONS

Anaerobic ciliates have been largely unaccounted for in our assessments of the diversity of anoxic microbial communities, despite their important ecological roles in biogeochemical cycling as grazers and via syntrophic associations with microbes from multiple domains of life. In this study, we leveraged publicly available 18S rDNA datasets, both from isolated, small-scale studies and larger-scale global sampling efforts, to explore anaerobic ciliate diversity across global marine oxygen-depleted habitats. Using a custom and now publicly available bioinformatics pipeline, we reprocessed 2854 environmental samples from diverse habitats and used phylogenetic placement methods to uncover a remarkable diversity of anaerobic ciliate sequences, including a substantial number which were divergent from known sequences and represent potentially novel lineages. For example, we found that the majority of anaerobic ciliate sequences belonged to the poorly described family Epalxellidae, highlighting the global ecological relevance and potential for discovery in this group. Additionally, we recovered many divergent sequences from the deep-branching clades Cariacotrichea, Parablepharismea, and Muranotrichea, which have rarely been described in culture-based studies.

Our results highlight the widespread presence of anaerobic ciliates across geographically and physically disparate marine habitats. We found that, in many cases, the habitats closest in community composition originated from shallow and deep-sea environments, potentially due to their similarities in redox structure and microbial community context, suggesting that anaerobic protists, and potentially other extremophile protists, are less structured by water depth and geography than other protist groups, at least at the taxonomic levels possible to differentiate with 18S rRNA amplicon data. Of course, it is possible that this data cannot differentiate species– or strain-level diversity, and that these habitats instead share similar community profiles at the genus, or some higher, taxonomic level. Future work with higher resolution sequencing approaches like full-length rRNA gene amplicons or population genomics will help to resolve these types of questions. For now, there is an abundance of publicly available short read sequencing data from diverse habitats around the globe which represent an underutilized resource in the study of poorly researched protist groups. This study demonstrates the utility of our pipeline for the recovery and discovery of short read sequences from any protist lineage, given a reference phylogenetic tree. Through our meta-analysis of publicly available sequencing data, we demonstrated that anaerobic ciliates are globally distributed members of marine protist communities with substantial reservoirs of uncharacterized diversity. Additionally, we provide a framework for other researchers to investigate the diversity and ecology of protists from other little-known taxonomic groups and/or under-sampled habitat types.

## ACKNOWLEDGEMENTS

The authors acknowledge support from the URI Center for Computational Research. In particular, the computations were performed on the UNITY high-performance computing cluster, a multi-institutional cluster led by University of Massachusetts Amherst, the University of Rhode Island, and University of Massachusetts Dartmouth. The authors would like to thank Dr. Johana Rotterova for her help with the curation of the APM reference phylogenetic tree, construction of Supplemental Table 3 (free-living anaerobic ciliate taxonomy), as well as sending along datasets to be included in the meta-analysis. We also thank Kateřina Koštířová for providing us with the reference alignment for the Anaerocyclidiidae reference phylogenetic tree and Dr. Cecile Cres for her help in debugging the Snakemake pipeline. Thank you also to Dr. Sarah Hu for providing a tutorial for using Snakemake with QIIME2, which this Snakemake pipeline was built off of: https://forum.qiime2.org/t/qiime2-snakemake-workflow-tutorial-18s-16s-tag-sequencing/11334. We thank Brenden Kelly for help with sediment core collection along with Stephen Granger and divers from the University of Rhode Island. This work was supported by a Simons Foundation Early Career Investigator in Marine Microbial Ecology and Evolution Award to RAB, the United States National Science Foundation EPSCoR Track II Cooperative Agreement Award #1330406, Rhode Island Seagrant, and the URI Coastal Institute. Sequencing was conducted at the Genomics and Sequencing Center, a RI NSF EPSCoR research facility. Sediment core collection was supported by RI Sea Grant funding to RF.

## AUTHOR CONTRIBUTION

AS and RAB conceptualized the work; AS completed the literature review and all analyses; RF and AH collected the Narragansett Bay, RI sediment samples; EF generated the 18S rDNA amplicons from the sediment samples; AS and RAB discussed and interpreted the results; RAB acquired funding and supervised the co-authors; AS wrote the manuscript with input and revisions from all co-authors.

## DATA AVAILABILITY

Raw 18S rDNA amplicon sequences which were generated in this study are available in the NCBI Sequence Read Archive (SRA) (Bioproject ID PRJNA1347111). Scripts for bioinformatic analyses, including the Snakemake pipeline, are available on GitHub, https://github.com/aschrecengost/18S_meta-analysis/tree/main.

## SUPPLEMENTAL INFORMATION

### METHODS

#### Study selection and metadata

The study selection criteria included: those which conducted Illumina amplicon sequencing of any region of the 18S rRNA gene, deposited the raw .fastq files in some public repository that could be accessed (usually NCBI SRA; in some cases, the corresponding authors were contacted for these files), and those that included samples from marine habitats which were or could possibly be oxygen-depleted, either explicitly with O_2_ measurements, in habitats known to be oxygen depleted (1), or that reported the presence of anaerobic ciliate taxa.

Metadata was obtained from the NCBI SRA website for each study and occasionally supplemented from the primary literature if necessary. The metadata was curated to include following fields for all studies (Table S2): Sample ID, Study ID, latitude, longitude, depth, primer region, primer set, sequencing chemistry, env_1 (indicates whether the sample is marine or brackish), env_2 (indicates whether the sample originated from sediment, water, or microbial mat), env_3 (indicates the habitat type e.g. intertidal, subtidal, hydrothermal vent), env_4 (the sample description as provided from the original metadata file/the literature), env_5 (our own most detailed description of habitat). Other metadata fields were also included for some samples when relevant, e.g. oxygen concentration measurements from the TARA oceans dataset to identify anoxic samples.

#### Sampling and sequencing of Narragansett Bay sediments

Sediment samples were collected from three sites across Narragansett Bay, RI, USA: the Providence River Estuary, Wickford Harbor (Public Dock and Gardener’s Wharf) and Mid Bay (Table S2). Sediment cores (10 cm inner diameter and 30.5 cm long) were collected using either a 4.5 m long pole corer (Providence River Estuary and Wickford Harbor) or with SCUBA divers (Mid Bay). With each method the vertical structure of the sediment core as well as the flocculent surface layer were retained (2). These cores were subsampled into multiple depth horizons (1 cm deep, up to 6 cm). DNA from the top two horizons from all cores, as well as the additional four depth horizons from the Wickford Harbor sites, was extracted from the sediment samples using the Macherey-Nagel Nucleospin Soil Kit (Macherey-Nagel, Düren, Germany), following manufacturer’s instructions. 18S rDNA was amplified from this DNA using a universal primer set targeting the V9 region of the 18S rRNA gene: 1389F (5′-TTGTACACACCGCCC-3′) and 1510R (5′-CCTTCYGCAGGTTCACCTAC-3′) (3), as well as a Stramenopile-Alveolata-Rhizaria (SAR) specific primer set: SAR F (5’-AYTCAGGGAGGTAGTGACAAG-3’) and SAR R (5’-RACTACGAGCTTTTTAACTGC-3’) (4). The resulting amplicons were sequenced on an Illumina MiSeq at the RI Genomics and Sequencing Center to generate paired end reads (2×300 bp).

#### SRA to ASV Snakemake pipeline: Raw data retrieval, processing into per-sample ASVs, and taxonomic assignment

Raw data retrieval and processing into ASVs was carried out via QIIME2 and Snakemake, with all steps implemented as modular Snakemake rules (Figure S1). Parameters were kept as consistent as possible to facilitate combinations of ASV tables, and when possible reads were trimmed to the exact same primer region. Raw FASTQ files from each study were retrieved from the NCBI Sequence Read Archive (SRA) via user input of SRA accession ID into the Snakefile. Run metadata was obtained via the Entrez Direct utilities esearch and efetch and filtered to extract valid SRR sample identifiers (5). FASTQ files were downloaded using parallel-fastq-dump, which supports multithreaded retrieval and automatic splitting of paired-end reads.

Raw FASTQ files were renamed to QIIME2 convention and then imported into QIIME2. Adapters and primers were removed with qiime cutadapt trim-paired with user-defined primer sequences and read-through trimming where necessary (6). Forward and reverse reads were merged using qiime vsearch merge-pairs, allowing staggered alignment with user-defined thresholds for mismatch tolerance and minimum overlap length (7). Merged reads were quality filtered with qiime quality-filter q-score, and denoising was performed using Deblur inside QIIME2 with reads trimmed to a user-defined fixed length and denoised against the Protist Ribosomal Reference Database (PR^2^) provided in FASTA format (8,9). This process generated for each study a FASTA file of ASVs, an ASV count table, and accompanying deblur statistics. These statistics along with the raw reads, primer-trimmed reads, merged reads, and filtered reads were visualized with qiime demux summarize on view.qiime2.org to diagnose potential issues during sequence processing.

Taxonomic assignment of representative sequences was performed using a Naive Bayes classifier trained on the PR^2^ reference database, implemented via qiime feature-classifier classify-sklearn (10). Classification output was used for downstream filtering: sequences classified under the phylum Ciliophora were filtered from both the feature table and representative sequence set using qiime taxa filter-table and qiime taxa filter-seqs, respectively, and the same was done for sequences which were classified under Unassigned at the kingdom or phylum level.

#### Generation of anaerobic ciliate clade phylogenetic trees

Full length 18S rRNA gene datasets were manually curated based on the literature and dereplicated at 99% identity with the aim of including one sequence for each species. For Plagiopylea, we included both cultured and environmental strains which have been published in phylogenies previously and used Prostomatea as an outgroup (11,12) (Figure S2). For Anaerocyclididiiae, we used a sequence alignment from a recently published study which established this family as a novel anaerobic lineage (13) (Figure S4). For the APM clade, we included both cultured and environmental sequences which have been included in published phylogenies of this group and also supplemented with environmental sequences from PR^2^ which were assigned to Cariacotrichea (14,15) (Figure S3). Full length 18S rRNA sequences were obtained from GenBank. Per-clade alignments were generated using the MAFFT algorithm and the progressive method L-INS-I (16). Alignments were manually trimmed using AliView (17) and phylogenetic trees were generated using a Maximum likelihood method in RaxML under the GTR+GAMMA substitution model with 200 independent runs and 1000 bootstrap replicates (18). Files which describe the taxonomy of each branch in the phylogeny were generated as well.

#### Analysis of phylogenetic placement results and anaerobic ciliate sequences

Gappa (19) was used to analyze and visualize phylogenetic placement results. The expected distance between placement locations (EDPL) and likelihood weight ratio (LWR) of queries and placements were calculated with gappa examine lwr-list and gappa examine edpl, respectively. Queries with EDPL > 0.05 were filtered out using a custom script and seqkit (20) and excluded from further analysis. In this way, sequences with diffuse and uncertain placements were removed, only retaining those placed with high confidence in a specific area of the tree. To translate phylogenetic placements into taxonomic annotations, we used the command gappa examine assign. This step maps the likelihood weight of each phylogenetic placement onto a user-specified taxonomy. The command was executed with the –-best-hit strategy, which assigns each query to the taxon associated with its highest-likelihood placement, and a consensus threshold of 0.7, which sets the minimum proportion of descendant tips that must share a taxonomic rank for the query to be assigned to an internal node.

To assess phylogenetic divergence between anaerobic ciliate communities from different habitat types, we performed another phylogenetic placement of all anaerobic ciliate ASVs onto a modified Ciliophora reference tree, which was re-inferred after the addition of all of the sequences from the anaerobic clade reference tree. Prior to analysis, .jplace files were split per sample using gappa edit split and the associated ASV count table. Only samples containing a minimum number of placements were retained for downstream analysis (at least 5). We grouped per-sample .jplace files by habitat type and used the command gappa analyze krd to compute the Kantorovich-Rubinstein Distance (KRD) between these grouped samples. The KRD is a generalization of Unifrac distance which allows read placements anywhere in the tree and allows for uncertainty in placement. It calculates the minimum effort needed to transform one community’s placement distribution into another’s by moving mass along the tree (21). The KRD matrix was converted to a distance object in R and hierarchically clustered using the UPGMA (Unweighted Pair Group Method with Arithmetic Mean) algorithm via the hclust() function.

We also performed squash clustering on these same groups of samples, which is a hierarchical clustering method designed for phylogenetic placement data. Squash clustering reduces each sample’s placements into a single averaged distribution in the tree, averages the placement profiles below each node in the tree, then measures the distance between these averaged distributions to define branch lengths (22). We applied squash clustering via the command gappa analyze squash to the same per-habitat .jplace files, and the tree topology was congruent with that which we obtained with KRD. Because uneven sampling depth and primer bias can compromise distance-based calculations, we were cautious about interpreting community divergence based on KRD alone. The observed clustering patterns from KRD were consistent with those obtained using squash clustering (Figure S5), which is more robust to differences in read depth because each sample is represented as a normalized placement distribution. The agreement between both methods suggests that our results reflect a biological signal rather than methodological bias.

For each anaerobic clade, we re-inferred phylogenetic trees by combining clade-assigned ASVs with reference sequences and aligning them using the MAFFT algorithm with the –-addfragments option, which is meant to add short sequences to a multiple sequence alignment, and maintaining the original alignment length (16). Maximum likelihood tree inference was performed with multithreaded RAxML v8.2 (18) under the GTR+GAMMA substitution model with 200 independent tree searches and 1000 bootstrap replicates. To preserve the reference phylogenetic tree topology, we used the per-clade reference trees as constraint trees with the –g parameter. Following tree inference, patristic distances, which is a type of phylogenetic distance defined as the sum of branch lengths connecting two nodes, were calculated between each query and reference sequence using the cophenetic.phylo() function from the ape package in R (23). For each query, the minimum patristic distance was extracted, yielding a per-query estimate of evolutionary divergence from the closest known relative. These distances were visualized using ridge plots in R to compare clade-specific patterns of phylogenetic novelty. Additionally, for each query sequence BLASTn searches were performed against the PR^2^ database to identify the percent identity of each ASV to its closest reference sequence (24). Percent identity values from the top BLAST hits were extracted and visualized using ridge plots in R.

As using the maximum reference patristic distance is the most conservative approach to categorize novel sequences, for each clade we provide a range of percentages of novel sequences, with the number calculated using the more conservative maximum threshold at the bottom of the range. Sequences which were assigned to Odontostomatida or whose closest neighbor was an odontostomatid were excluded from this classification, because the poorly resolved phylogenetic position of this group renders calculated phylogenetic distances unreliable (25).

## SUPPLEMENTAL TABLES

**Table S1.** Studies included in meta-analysis.

**Table S2.** Sample metadata.

**Table S3.** Known free-living anaerobic ciliate taxa.

**Table S4.** Classification of novel ASVs based on patristic distances.

**Supplemental Figure 1.**
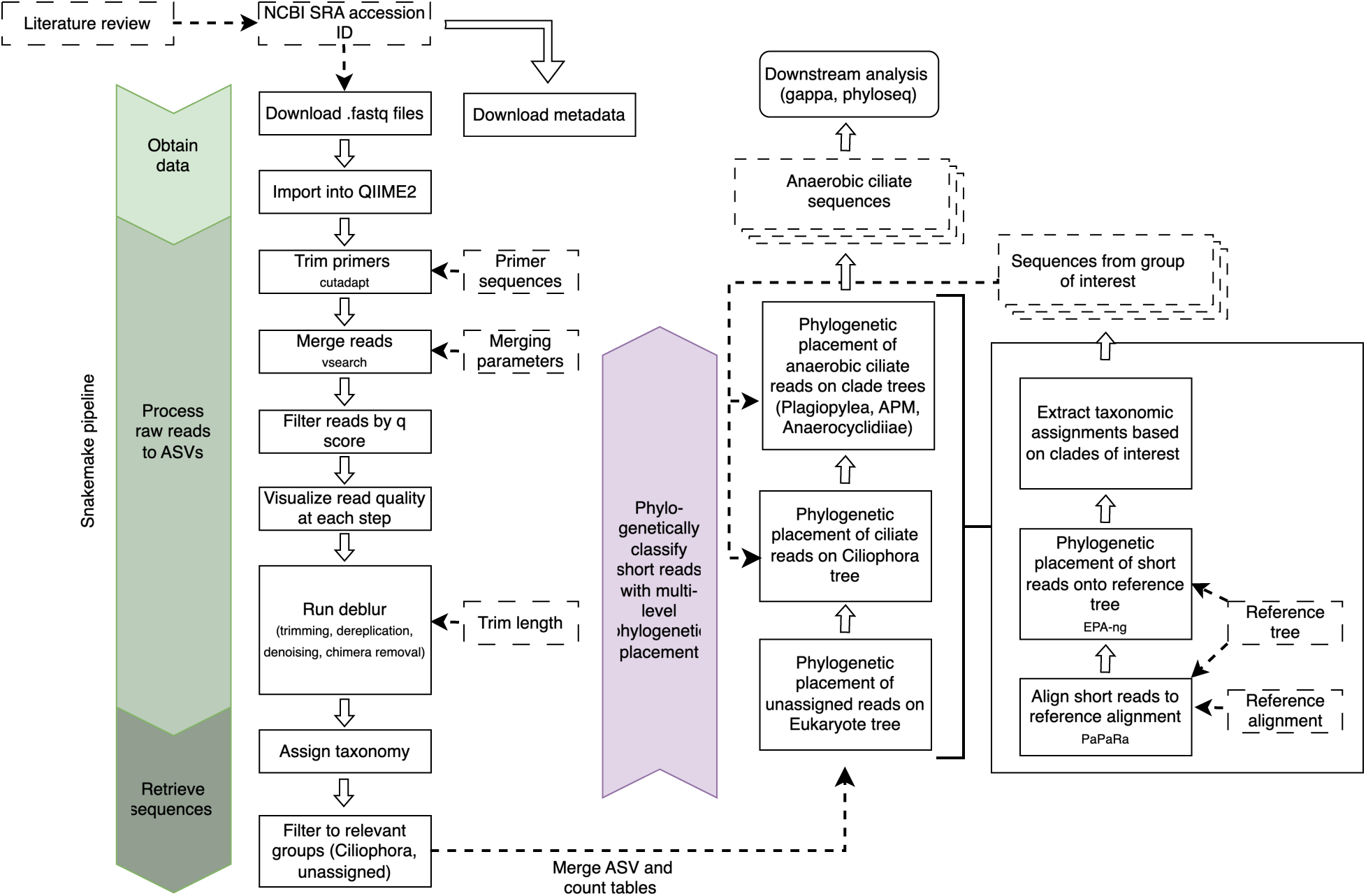
Flowchart describing methodology, from literature review to obtaining anaerobic ciliate sequences. Snakemake pipeline is shown at left in green. User-supplied inputs are represented by dashed-line boxes.

**Supplemental Figure 2.**
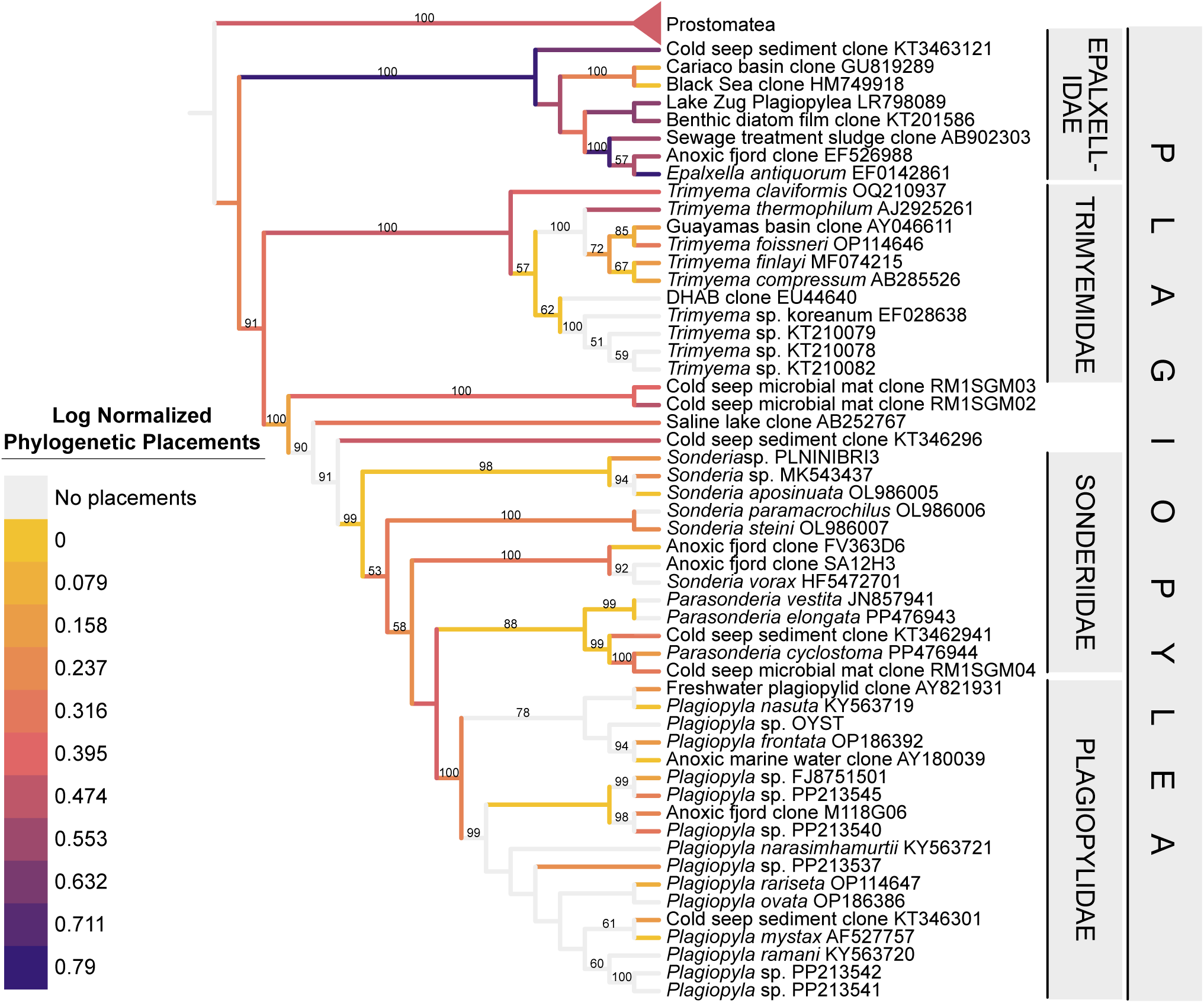
Phylogenetic placement results of plagiopylid ciliates. Queries which were assigned to Plagiopylea after being placed on the Ciliophora reference tree were then placed onto the Plagiopylea reference tree, which is pictured here. Bootstrap values on reference tree branches are based on 1000 replicates and only values greater than 50 are included. Color on branches indicates log-normalized distribution of top phylogenetic placements for each query, which corresponds to their phylogenetic-placement based taxonomic assignment. Ciliate family is labelled on the right of the tree.

**Supplemental Figure 3.**
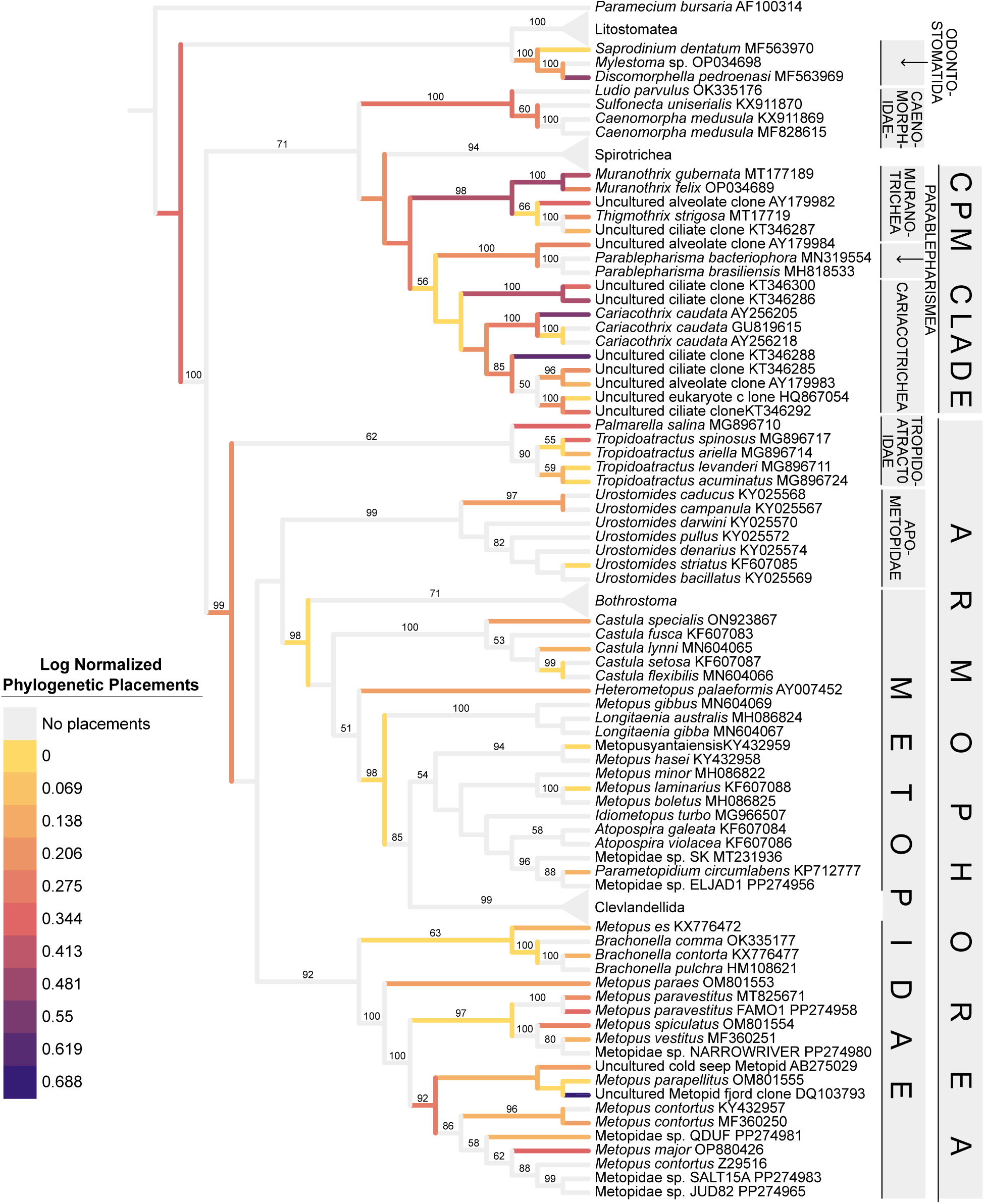
Phylogenetic placement results of APM ciliates. Queries which were assigned to the APM clade after being placed on the Ciliophora reference tree were then placed onto the APM reference tree, which is pictured here. Color on branches indicates log-normalized distribution of top phylogenetic placements for each query, which corresponds to their phylogenetic-placement based taxonomic assignment. Bootstrap values on reference tree branches are based on 1000 replicates and only values greater than 50 are included. Ciliate family/order/class is labelled on the right of the tree.

**Supplemental Figure 4.**
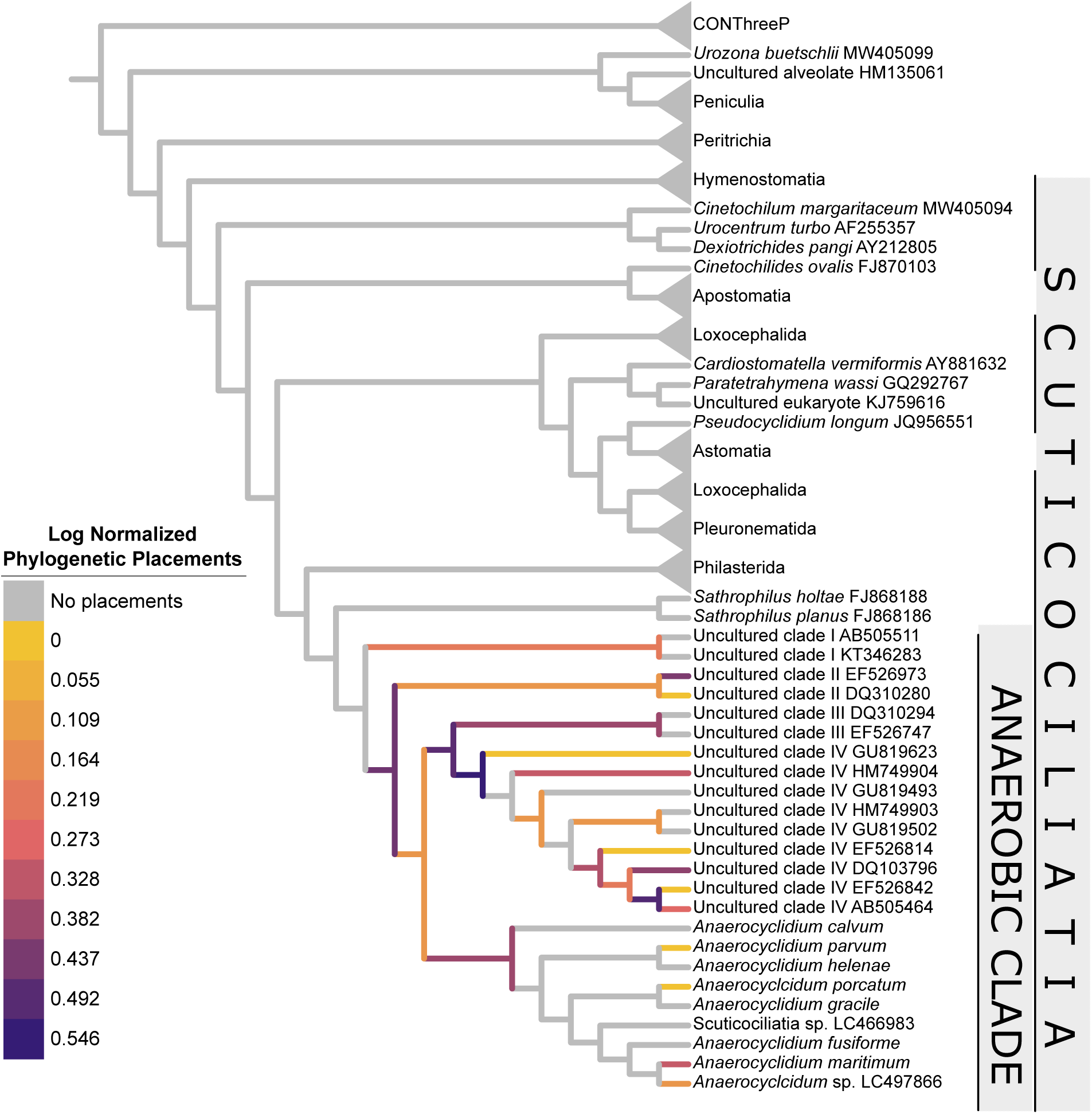
Phylogenetic placement results of Anaerocyclidiidae ciliates. Queries which were assigned to the Anaerocyclidiidae clade after being placed on the Ciliophora reference tree were then placed onto the Anaerocyclidiidae reference tree, which is pictured here. Color on branches indicates log-normalized distribution of top phylogenetic placements for each query, which corresponds to their phylogenetic-placement based taxonomic assignment. Ciliate family/class is labelled on the right of the tree.

**Supplemental Figure 5.**
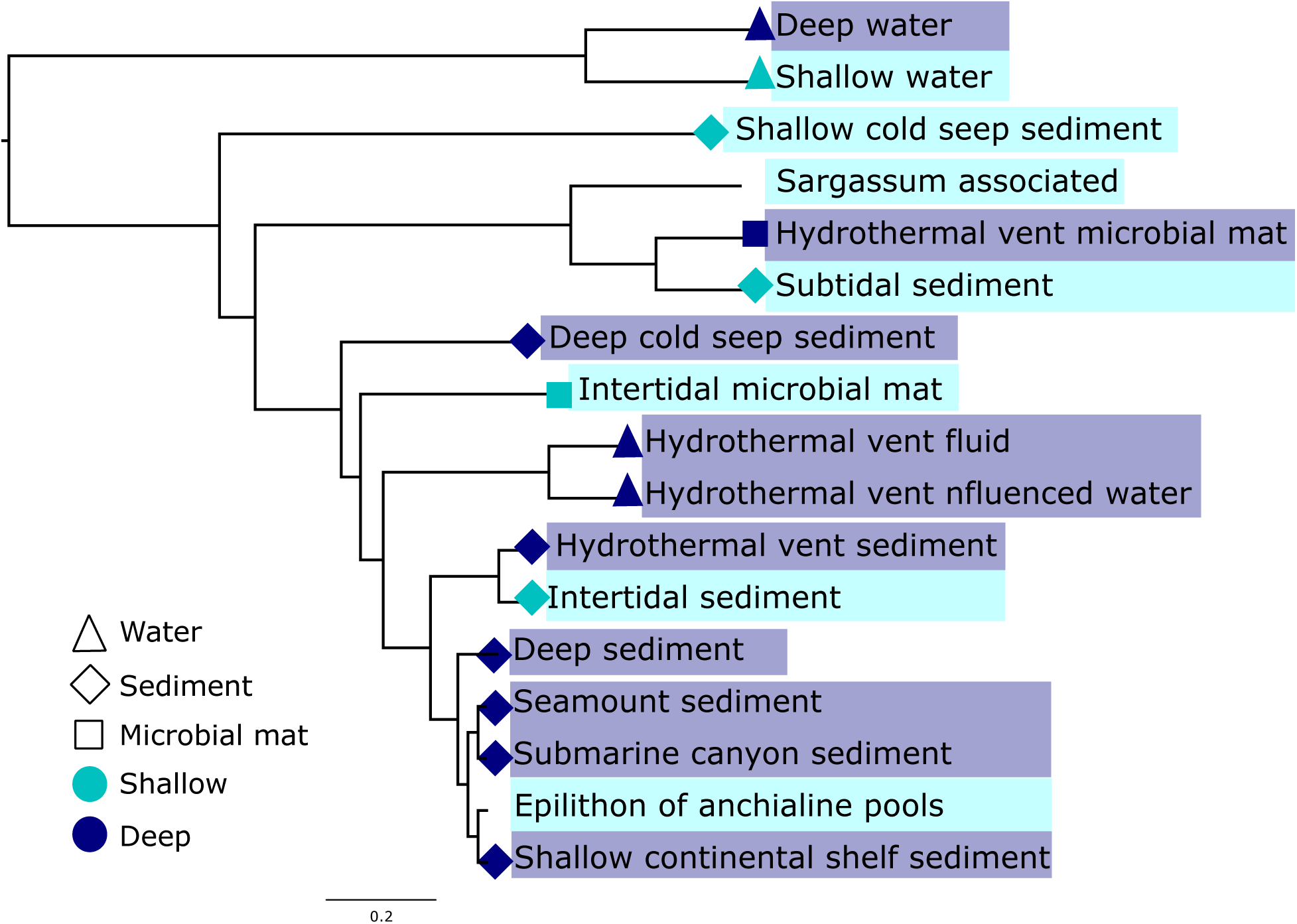
Squash clustering of samples grouped by habitat type. Branch tips are colored by water depth, with deep defined as greater than 300 m, and branch node shapes indicate sample substrate.

## REFERENCES

1. de Vargas C, Audic S, Henry N, Decelle J, Mahe F, Logares R, et al. Eukaryotic plankton diversity in the sunlit ocean. Science. 2015 May 22;348(6237):1261605–1261605.

2. DeLong EF, Karl DM. Genomic perspectives in microbial oceanography. Nature. 2005 Sept;437(7057):336–42.

3. Caron DA, Worden AZ, Countway PD, Demir E, Heidelberg KB. Protists are microbes too: a perspective. ISME J. 2009 Jan;3(1):4–12.

4. Vaulot D, Sim CWH, Ong D, Teo B, Biwer C, Jamy M, et al. metaPR2: a database of eukaryotic 18S rRNA metabarcodes with an emphasis on protists. Mol Ecol Resour. 2022;22(8):3188–201.

5. Forster D, Dunthorn M, Mahé F, Dolan JR, Audic S, Bass D, et al. Benthic protists: the under-charted majority. FEMS Microbiol Ecol. 2016;92(8).

6. Sarmiento JL, Gruber N. Ocean Biogeochemical dynamics. Princeton University Press; 2006.

7. Cavicchioli R, Ripple WJ, Timmis KN, Azam F, Bakken LR, Baylis M, et al. Scientists’ warning to humanity: microorganisms and climate change. Nat Rev Microbiol. 2019;17(9):569–86.

8. Santoferrara LF. Current practice in plankton metabarcoding: optimization and error management. J Plankton Res. 2019;41(5):571–82.

9. Burki F, Sandin MM, Jamy M. Diversity and ecology of protists revealed by metabarcoding. Curr Biol. 2021 Oct 11;31(19):R1267–80.

10. Kopf A, Bicak M, Kottmann R, Schnetzer J, Kostadinov I, Lehmann K, et al. The ocean sampling day consortium. GigaScience. 2015 Dec 1;4(1):s13742-015-0066–5.

11. Duarte CM. Seafaring in the 21st century: the Malaspina 2010 circumnavigation expedition. 2015.

12. Fenchel T, Finlay BJ. The biology of free-living anaerobic ciliates. Eur J Protistol. 1991;26(3– 4):201–15.

13. Dziallas C, Allgaier M, Monaghan MT, Grossart HP. Act together—implications of symbioses in aquatic ciliates. Front Microbiol. 2012 Aug 7;3.

14. Fokin SI. Frequency and biodiversity of symbionts in representatives of the main classes of Ciliophora. Eur J Protistol. 2012 May 1;48(2):138–48.

15. Rotterová J, Salomaki E, Pánek T, Bourland W, Žihala D, Táborskỳ P, et al. Genomics of New Ciliate Lineages Provides Insight into the Evolution of Obligate Anaerobiosis. Curr. Biol. 2020 Jun 8;30(11):2037–50.

16. Fenchel T, Finlay BJ. Free-living protozoa with endosymbiotic methanogens. In: (Endo) symbiotic methanogenic archaea. Springer; 2010. p. 1–11.

17. Stoeck T, Epstein S. Novel Eukaryotic Lineages Inferred from Small-Subunit rRNA Analyses of Oxygen-Depleted Marine Environments. Appl Environ Microbiol. 2003 May;69(5):2657– 63.

18. Bernhard JM, Buck KR, Farmer MA, Bowser SS. The Santa Barbara Basin is a symbiosis oasis. Nature. 2000;403(6765):77–80.

19. Fenchel T, Finlay BJ. The Evolution of Life without Oxygen. Am Sci. 1994;82(1):22–9.

20. Rotterová J, Edgcomb VP, Čepička I, Beinart R. Anaerobic Ciliates as a Model Group for Studying Symbioses in Oxygen-depleted Environments. J Eukaryot Microbiol. 2022 Sep;69(5):e12912.

21. Fenchel T. Anaerobic Eukaryotes. In: Altenbach AV, Bernhard JM, Seckbach J, editors. Anoxia. Dordrecht: Springer Netherlands; 2012. p. 3–16. (Cellular Origin, Life in Extreme Habitats and Astrobiology; vol. 21).

22. van Hoek AHAM, van Alen TA, Sprakel VSI, Leunissen JAM, Brigge T, Vogels GD, et al. Multiple Acquisition of Methanogenic Archaeal Symbionts by Anaerobic Ciliates. Mol Biol Evol. 2000 Feb 1;17(2):251–8.

23. Stairs CW, Leger MM, Roger AJ. Diversity and origins of anaerobic metabolism in mitochondria and related organelles. Philos Trans R Soc B Biol Sci. 2015;370(1678):20140326.

24. Pester M, Knorr KH, Friedrich MW, Wagner M, Loy A. Sulfate-reducing microorganisms in wetlands–fameless actors in carbon cycling and climate change. Front Microbiol. 2012;3:72.

25. Worden AZ, Follows MJ, Giovannoni SJ, Wilken S, Zimmerman AE, Keeling PJ. Rethinking the marine carbon cycle: factoring in the multifarious lifestyles of microbes. Science. 2015;347(6223):1257594.

26. Diaz RJ, Rosenberg R. Spreading dead zones and consequences for marine ecosystems. science. 2008;321(5891):926–9.

27. Schmidtko S, Stramma L, Visbeck M. Decline in global oceanic oxygen content during the past five decades. Nature. 2017 Feb;542(7641):335–9.

28. Warren A, Patterson DJ, Dunthorn M, Clamp JC, Achilles-Day UEM, Aescht E, et al. Beyond the “Code”: A Guide to the Description and Documentation of Biodiversity in Ciliated Protists (Alveolata, Ciliophora). J Eukaryot Microbiol. 2017 July 1;64(4):539–54.

29. López-García P, Philippe H, Gail F, Moreira D. Autochthonous eukaryotic diversity in hydrothermal sediment and experimental microcolonizers at the Mid-Atlantic Ridge. Proc Natl Acad Sci. 2003;100(2):697–702.

30. Takishita K, Miyake H, Kawato M, Maruyama T. Genetic diversity of microbial eukaryotes in anoxic sediment around fumaroles on a submarine caldera floor based on the small-subunit rDNA phylogeny. Extremophiles. 2005 June;9(3):185–96.

31. Dawson SC, Pace NR. Novel kingdom-level eukaryotic diversity in anoxic environments. Proc Natl Acad Sci. 2002 June 11;99(12):8324–9.

32. Behnke A, Bunge J, Barger K, Breiner HW, Alla V, Stoeck T. Microeukaryote community patterns along an O2/H2S gradient in a supersulfidic anoxic fjord (Framvaren, Norway). Appl Environ Microbiol. 2006;72(5):3626–36.

33. Edgcomb VP, Kysela DT, Teske A, de Vera Gomez A, Sogin ML. Benthic eukaryotic diversity in the Guaymas Basin hydrothermal vent environment. Proc Natl Acad Sci. 2002;99(11):7658– 62.

34. Takishita K, Kakizoe N, Yoshida T, Maruyama T. Molecular evidence that phylogenetically diverged ciliates are active in microbial mats of deep-sea cold-seep sediment. J Eukaryot Microbiol. 2010;57(1):76–86.

35. Orsi W, Edgcomb V, Faria J, Foissner W, Fowle WH, Hohmann T, et al. Class Cariacotrichea, a novel ciliate taxon from the anoxic Cariaco Basin, Venezuela. Int J Syst Evol Microbiol. 62(Pt_6):1425–33.

36. Stoeck T, Bass D, Nebel M, Christen R, Jones MDM, Breiner HW, et al. Multiple marker parallel tag environmental DNA sequencing reveals a highly complex eukaryotic community in marine anoxic water. Mol Ecol. 2010;19(s1):21–31.

37. Edgcomb V, Orsi W, Bunge J, Jeon S, Christen R, Leslin C, et al. Protistan microbial observatory in the Cariaco Basin, Caribbean. I. Pyrosequencing vs Sanger insights into species richness. ISME J. 2011;5(8):1344–56.

38. Stoeck T, Behnke A, Christen R, Amaral-Zettler L, Rodriguez-Mora MJ, Chistoserdov A, et al. Massively parallel tag sequencing reveals the complexity of anaerobic marine protistan communities. BMC Biol. 2009 Nov 3;7(1):72.

39. Salonen IS, Chronopoulou PM, Leskinen E, Koho KA. Metabarcoding successfully tracks temporal changes in eukaryotic communities in coastal sediments. FEMS Microbiol Ecol. 2019 Jan 1;95(1):fiy226.

40. Zhao F, Xu K. Distribution of Ciliates in Intertidal Sediments across Geographic Distances: A Molecular View. Protist. 2017 Apr 1;168(2):171–82.

41. Kong J, Wang Y, Warren A, Huang B, Sun P. Diversity Distribution and Assembly Mechanisms of Planktonic and Benthic Microeukaryote Communities in Intertidal Zones of Southeast Fujian, China. Front Microbiol. 2019 Nov 15;10:2640

42. Murdock SA, Juniper SK. Hydrothermal vent protistan distribution along the Mariana arc suggests vent endemics may be rare and novel. Environ Microbiol. 2019 Oct;21(10):3796– 815.

43. Pasulka A, Hu SK, Countway PD, Coyne KJ, Cary SC, Heidelberg KB, et al. SSU-rRNA Gene Sequencing Survey of Benthic Microbial Eukaryotes from Guaymas Basin Hydrothermal Vent. J Eukaryot Microbiol. 2019;66(4):637–53.

44. Pasulka AL, Levin LA, Steele JA, Case DH, Landry MR, Orphan VJ. Microbial eukaryotic distributions and diversity patterns in a deep-sea methane seep ecosystem. Environ Microbiol. 2016;18(9):3022–43.

45. Zhao F, Filker S, Wang C, Xu K. Bathymetric gradient shapes the community composition rather than the species richness of deep-sea benthic ciliates. Sci Total Environ. 2021 Feb 10;755:142623.

46. Hu SK, Smith AR, Anderson RE, Sylva SP, Setzer M, Steadmon M, et al. Globally-distributed microbial eukaryotes exhibit endemism at deep-sea hydrothermal vents. Mol Ecol. 2022;

47. Mars Brisbin M, Conover AE, Mitarai S. Influence of regional oceanography and hydrothermal activity on protist diversity and community structure in the Okinawa Trough. Microb Ecol. 2020;80:746–61.

48. Schrecengost A, Rotterová J, Poláková K, Čepička I, Beinart RA. Divergent marine anaerobic ciliates harbor closely related Methanocorpusculum endosymbionts. ISME J. 2024 Jan 1;18(1):wrae125.

49. Méndez-Sánchez D, Schrecengost A, Rotterová J, Koštířová K, Beinart RA, Čepička I. Methanogenic symbionts of anaerobic ciliates are host and habitat specific. ISME J. 2024;18(1):wrae164.

50. Li R, Zhuang W, Feng X, Al-Farraj SA, Schrecengost A, Rotterova J, et al. Molecular phylogeny and taxonomy of three anaerobic plagiopyleans (Alveolata: Ciliophora), retrieved from two geographically distant localities in Asia and North America. Zool J Linn Soc. 2023;zlad015.

51. Czech L, Barbera P, Stamatakis A. Methods for automatic reference trees and multilevel phylogenetic placement. Bioinformatics. 2019 Apr 1;35(7):1151–8.

52. Mahé F, de Vargas C, Bass D, Czech L, Stamatakis A, Lara E, et al. Parasites dominate hyperdiverse soil protist communities in Neotropical rainforests. Nat Ecol Evol. 2017;1(4):0091.

53. Nitla V, Serra V, Fokin SI, Modeo L, Verni F, Sandeep BV, et al. Critical revision of the family Plagiopylidae (Ciliophora: Plagiopylea), including the description of two novel species, Plagiopyla ramani and Plagiopyla narasimhamurtii, and redescription of Plagiopyla nasuta Stein, 1860 from India. Zool J Linn Soc. 2019 Apr 22;186(1):1–45.

54. Poláková K, Bourland WA, Čepička I. Anaerocyclidiidae fam. nov.(Oligohymenophorea, Scuticociliatia): a newly recognized major lineage of anaerobic ciliates hosting prokaryotic symbionts. Eur J Protistol. 2023;90:126009.

55. Ewers I, Rajter L, Czech L, Mahé F, Stamatakis A, Dunthorn M. Interpreting phylogenetic placements for taxonomic assignment of environmental DNA. J Eukaryot Microbiol. 2023;70(5):e12990.

56. Czech L, Stamatakis A, Dunthorn M, Barbera P. Metagenomic Analysis Using Phylogenetic Placement—A Review of the First Decade. Front Bioinformatics. 2022 May 26;2:871393.

57. Stoeck T, Foissner W, Lynn DH. Small-Subunit rRNA Phylogenies Suggest That Epalxella antiquorum (Penard, 1922) Corliss, 1960 (Ciliophora, Odontostomatida) Is a Member of the Plagyopylea. J Eukaryot Microbiol. 2007;54(5):436–42.

58. Fernandes NM, Vizzoni VF, Borges B do N, Soares CAG, da Silva-Neto ID, Paiva T da S. Molecular phylogeny and comparative morphology indicate that odontostomatids (Alveolata, Ciliophora) form a distinct class-level taxon related to Armophorea. Mol Phylogenet Evol. 2018 Sept;126:382–9.

59. Bolyen E, Rideout JR, Dillon MR, Bokulich NA, Abnet CC, Al-Ghalith GA, et al. Reproducible, interactive, scalable and extensible microbiome data science using QIIME 2. Nat Biotechnol. 2019 Aug;37(8):852–7.

60. Amir A, McDonald D, Navas-Molina JA, Kopylova E, Morton JT, Zech Xu Z, et al. Deblur Rapidly Resolves Single-Nucleotide Community Sequence Patterns. mSystems. 2(2):e00191–16.

61. Rognes T, Flouri T, Nichols B, Quince C, Mahé F. VSEARCH: a versatile open source tool for metagenomics. PeerJ. 2016 Oct 18;4:e2584.

62. Martin M. Cutadapt removes adapter sequences from high-throughput sequencing reads. EMBnet.journal. 2011 May 2;17(1):10–2.

63. Bokulich NA, Kaehler BD, Rideout JR, Dillon M, Bolyen E, Knight R, et al. Optimizing taxonomic classification of marker-gene amplicon sequences with QIIME 2’s q2-feature-classifier plugin. Microbiome. 2018 May 17;6(1):90.

64. Rajter Ľ, Dunthorn M. Ciliate SSU-rDNA reference alignments and trees for phylogenetic placements of metabarcoding data. Metabarcoding Metagenomics. 2021 Aug 30;5:e69602.

65. Berger SA, Stamatakis A. Aligning short reads to reference alignments and trees. Bioinformatics. 2011;27(15):2068–75.

66. Barbera P, Kozlov AM, Czech L, Morel B, Darriba D, Flouri T, et al. EPA-ng: Massively Parallel Evolutionary Placement of Genetic Sequences. Syst Biol. 2019 Mar 1;68(2):365–9.

67. Kozlov AM, Darriba D, Flouri T, Morel B, Stamatakis A. RAxML-NG: a fast, scalable and user-friendly tool for maximum likelihood phylogenetic inference. Bioinformatics. 2019 Nov 1;35(21):4453–5.

68. Czech L, Barbera P, Stamatakis A. Genesis and Gappa: processing, analyzing and visualizing phylogenetic (placement) data. Bioinformatics. 2020 May 15;36(10):3263–5.

69. Letunic I, Bork P. Interactive Tree of Life (iTOL) v6: recent updates to the phylogenetic tree display and annotation tool. Nucleic Acids Res. 2024 July 5;52(W1):W78–82.

70. Pebesma E, Bivand R. Spatial data science: With applications in R. Chapman and Hall/CRC; 2023 May 10.

71. Villanueva RAM, Chen ZJ. ggplot2: Elegant Graphics for Data Analysis (2nd ed.). Meas Interdiscip Res Perspect. 2019 July 3;17(3):160–7.

72. McMurdie PJ, Holmes S. phyloseq: An R Package for Reproducible Interactive Analysis and Graphics of Microbiome Census Data. PLOS ONE. 2013 Apr 22;8(4):e61217.

73. Team RC. R: A language and environment for statistical computing. Published online 2020. 2021.

74. Evans SN, Matsen FA. The phylogenetic Kantorovich–Rubinstein metric for environmental sequence samples. J R Stat Soc Ser B Stat Methodol. 2012;74(3):569–92.

75. Matsen IV FA, Evans SN. Edge principal components and squash clustering: using the special structure of phylogenetic placement data for sample comparison. PloS One. 2013;8(3):e56859.

76. Bianchi D, Weber TS, Kiko R, Deutsch C. Global niche of marine anaerobic metabolisms expanded by particle microenvironments. Nat Geosci. 2018 Apr;11(4):263–8.

77. Byrne N, Strous M, Crépeau V, Kartal B, Birrien JL, Schmid M, et al. Presence and activity of anaerobic ammonium-oxidizing bacteria at deep-sea hydrothermal vents. ISME J. 2009 Jan 1;3(1):117–23.

78. Liu F, Ding J, Zeng J, Wang C, Wu B, Yan Q, et al. Mangrove sediments are environmental hotspots for pathogenic protists. J Hazard Mater. 2024 Apr 5;467:133643.

79. Vieillard AM, Newell SE, Thrush SF. Recovering From Bias: A Call for Further Study of Underrepresented Tropical and Low-Nutrient Estuaries. J Geophys Res Biogeosciences. 2020;125(7):e2020JG005766.

80. Gloor GB, Macklaim JM, Pawlowsky-Glahn V, Egozcue JJ. Microbiome datasets are compositional: and this is not optional. Front Microbiol. 2017;8:2224.

81. Weiss J, Esteban GF. Tracking down the rare ciliate biosphere. Front Protistol. 2024 Jan 8;1:1308546.

82. Reveillaud J, Reddington E, McDermott J, Algar C, Meyer JL, Sylva S, et al. Subseafloor microbial communities in hydrogen-rich vent fluids from hydrothermal systems along the Mid-Cayman Rise. Environ Microbiol. 2016;18(6):1970–87.

83. Kirchman DL. Processes in anoxic environments. Process Microb Ecol. 2012;92–112.

34. Gasol JM (Editor). Microbial Ecology of the Oceans. John Wiley & Sons; 2018 Mar 27.

85. Bolhuis H, Cretoiu MS, Stal LJ. Molecular ecology of microbial mats. FEMS Microbiol Ecol. 2014;90(2):335–50.

86. Jean MRN, Gonzalez-Rizzo S, Gauffre-Autelin P, Lengger SK, Schouten S, Gros O. Two New Beggiatoa Species Inhabiting Marine Mangrove Sediments in the Caribbean. PLoS ONE. 2015 Feb 17;10(2):e0117832.

87. Hu SK, Herrera EL, Smith AR, Pachiadaki MG, Edgcomb VP, Sylva SP, et al. Protistan grazing impacts microbial communities and carbon cycling at deep-sea hydrothermal vents. bioRxiv. 2021 Feb 8;2021.02.08.430233.

88. Schnetzer A, Moorthi SD, Countway PD, Gast RJ, Gilg IC, Caron DA. Depth matters: Microbial eukaryote diversity and community structure in the eastern North Pacific revealed through environmental gene libraries. Deep Sea Res Part Oceanogr Res Pap. 2011 Jan 1;58(1):16–26.

89. Ollison GA, Hu SK, Mesrop LY, DeLong EF, Caron DA. Come rain or shine: Depth not season shapes the active protistan community at station ALOHA in the North Pacific Subtropical Gyre. Deep Sea Res Part Oceanogr Res Pap. 2021 Apr 1;170:103494.

90. Dünn M, Arndt H. Distribution Patterns of Benthic Protist Communities Depending on Depth Revealed by Environmental Sequencing—From the Sublittoral to the Deep Sea. Microorganisms. 2023 July;11(7):1664.

91. Gong J, Shi F, Ma B, Dong J, Pachiadaki M, Zhang X, et al. Depth shapes α– and β-diversities of microbial eukaryotes in surficial sediments of coastal ecosystems. Environ Microbiol. 2015;17(10):3722–37.

92. Gimmler A, Korn R, de Vargas C, Audic S, Stoeck T. The Tara Oceans voyage reveals global diversity and distribution patterns of marine planktonic ciliates. Sci Rep. 2016 Sept 16;6(1):33555.

93. Scheckenbach F, Hausmann K, Wylezich C, Weitere M, Arndt H. Large-scale patterns in biodiversity of microbial eukaryotes from the abyssal sea floor. Proc Natl Acad Sci. 2010 Jan 5;107(1):115–20.

94. Graf JS, Schorn S, Kitzinger K, Ahmerkamp S, Woehle C, Huettel B, et al. Anaerobic endosymbiont generates energy for ciliate host by denitrification. Nature. 2021 Mar;591(7850):445–50.

95. Orsi W, Edgcomb V, Faria J, Foissner W, Fowle WH, Hohmann T, et al. Class Cariacotrichea, a novel ciliate taxon from the anoxic Cariaco Basin, Venezuela. Int J Syst Evol Microbiol. 2012 June 1;62(Pt_6):1425–33.

96. Stoeck T, Taylor GT, Epstein SS. Novel eukaryotes from the permanently anoxic Cariaco Basin (Caribbean Sea). Appl Environ Microbiol. 2003 Sept;69(9):5656–63.

97. Boscaro V, Santoferrara LF, Zhang Q, Gentekaki E, Syberg-Olsen MJ, Del Campo J, et al. EukRef-Ciliophora: a manually curated, phylogeny-based database of small subunit rRNA gene sequences of ciliates. Environ Microbiol. 2018 June;20(6):2218–30.

98. Orsi W, Song YC, Hallam S, Edgcomb V. Effect of oxygen minimum zone formation on communities of marine protists. ISME J. 2012 Aug;6(8):1586–601.

99. Campello-Nunes PH, Fernandes NM, Szokoli F, Fokin SI, Serra V, Modeo L, et al. Parablepharisma (Ciliophora) is not a Heterotrich: A Phylogenetic and Morphological Study with the Proposal of New Taxa. Protist. 2020 Feb 7;125716.

## REFERENCES

1. Fenchel T. and Finlay B. J. Ecology and Evolution in Anoxic Worlds. Oxford University Press; 1995 Mar 16.

2. Fulweiler RW, Nixon SW. Net sediment N2 fluxes in a southern New England estuary: variations in space and time. Biogeochemistry. 2012 Nov;111(1–3):111–24.

3. Amaral-Zettler LA, McCliment EA, Ducklow HW, Huse SM. A method for studying protistan diversity using massively parallel sequencing of V9 hypervariable regions of small-subunit ribosomal RNA genes. PloS One. 2009;4(7):e6372.

4. Sisson C, Gulla-Devaney B, Katz LA, Grattepanche JD. Seed bank and seasonal patterns of the eukaryotic SAR (Stramenopila, Alveolata and Rhizaria) clade in a New England vernal pool. J Plankton Res. 2018 July 1;40(4):376–90.

5. Kans J. Entrez Direct: E-utilities on the Unix Command Line. In: Entrez Programming Utilities Help [Internet] 2024 Dec 23. National Center for Biotechnology Information (US).

6. Martin M. Cutadapt removes adapter sequences from high-throughput sequencing reads. EMBnet.journal. 2011 May 2;17(1):10–2.

7. Rognes T, Flouri T, Nichols B, Quince C, Mahé F. VSEARCH: a versatile open source tool for metagenomics. PeerJ. 2016 Oct 18;4:e2584.

8. Amir A, McDonald D, Navas-Molina JA, Kopylova E, Morton JT, Zech Xu Z, et al. Deblur Rapidly Resolves Single-Nucleotide Community Sequence Patterns. mSystems. 2(2):e00191–16.

9. Guillou L, Bachar D, Audic S, Bass D, Berney C, Bittner L, et al. The Protist Ribosomal Reference database (PR2): a catalog of unicellular eukaryote Small Sub-Unit rRNA sequences with curated taxonomy. Nucleic Acids Res. 2013 Jan;41(Database issue):D597– 604.

10. Bokulich NA, Kaehler BD, Rideout JR, Dillon M, Bolyen E, Knight R, et al. Optimizing taxonomic classification of marker-gene amplicon sequences with QIIME 2’s q2-feature-classifier plugin. Microbiome. 2018 May 17;6(1):90.

11. Nitla V, Serra V, Fokin SI, Modeo L, Verni F, Sandeep BV, et al. Critical revision of the family Plagiopylidae (Ciliophora: Plagiopylea), including the description of two novel species, Plagiopyla ramani and Plagiopyla narasimhamurtii, and redescription of Plagiopyla nasuta Stein, 1860 from India. Zool J Linn Soc. 2019 Apr 22;186(1):1–45.

12. Li R, Zhuang W, Feng X, Al-Farraj SA, Schrecengost A, Rotterova J, et al. Molecular phylogeny and taxonomy of three anaerobic plagiopyleans (Alveolata: Ciliophora), retrieved from two geographically distant localities in Asia and North America. Zool J Linn Soc. 2023;zlad015.

13. 1Poláková K, Bourland WA, Čepička I. Anaerocyclidiidae fam. nov.(Oligohymenophorea, Scuticociliatia): a newly recognized major lineage of anaerobic ciliates hosting prokaryotic symbionts. Eur J Protistol. 2023;90:126009.

14. Rotterová J, Salomaki E, Pánek T, Bourland W, Žihala D, Táborskỳ P, et al. Genomics of New Ciliate Lineages Provides Insight into the Evolution of Obligate Anaerobiosis. Curr Biol. 2020 Jun 8;30(11):2037–50.

15. Bourland W, Pomahač O, Čepička I. Morphology and phylogeny of two anaerobic freshwater ciliates: Brachonella comma sp. nov. and the widely distributed but little-known caenomorphid, Ludio parvulus Penard, 1922. J Eukaryot Microbiol. 2022 May;69(3):e12892.

16. Katoh K, Rozewicki J, Yamada KD. MAFFT online service: multiple sequence alignment, interactive sequence choice and visualization. Brief Bioinform. 2019 July 19;20(4):1160– 6.

17. Larsson A. AliView: a fast and lightweight alignment viewer and editor for large datasets. Bioinformatics. 2014;30(22):3276–8.

18. Stamatakis A. RAxML version 8: a tool for phylogenetic analysis and post-analysis of large phylogenies. Bioinformatics. 2014;30(9):1312–3.

19. Czech L, Barbera P, Stamatakis A. Genesis and Gappa: processing, analyzing and visualizing phylogenetic (placement) data. Bioinformatics. 2020 May 15;36(10):3263–5.

20. Shen W, Le S, Li Y, Hu F. SeqKit: a cross-platform and ultrafast toolkit for FASTA/Q file manipulation. PloS One. 2016;11(10):e0163962.

21. vans SN, Matsen FA. The phylogenetic Kantorovich–Rubinstein metric for environmental sequence samples. J R Stat Soc Ser B Stat Methodol. 2012;74(3):569–92.

22. Matsen IV FA, Evans SN. Edge principal components and squash clustering: using the special structure of phylogenetic placement data for sample comparison. PloS One. 2013;8(3):e56859.

23. Paradis E, Schliep K. ape 5.0: an environment for modern phylogenetics and evolutionary analyses in R. Bioinformatics. 2019;35(3):526–8.

24. Camacho C, Coulouris G, Avagyan V, Ma N, Papadopoulos J, Bealer K, et al. BLAST+: architecture and applications. BMC Bioinformatics. 2009 Dec;10(1):421.

25. Fernandes NM, Vizzoni VF, Borges B do N, Soares CAG, da Silva-Neto ID, Paiva T da S. Molecular phylogeny and comparative morphology indicate that odontostomatids (Alveolata, Ciliophora) form a distinct class-level taxon related to Armophorea. Mol Phylogenet Evol. 2018 Sept;126:382–9.

